# An EDS1-SAG101 complex functions in TNL-mediated immunity in *Solanaceae*

**DOI:** 10.1101/511956

**Authors:** Johannes Gantner, Jana Ordon, Carola Kretschmer, Raphaël Guerois, Johannes Stuttmann

## Abstract

EDS1 (Enhanced disease susceptibility 1) forms mutually exclusive heterodimers with its interaction partners PAD4 (Phytoalexin-deficient 4) and SAG101 (Sensecence-associated gene 101). Collectively, these complexes are required for resistance responses mediated by nucleotide-binding leucine-rich repeat-type immune receptors (NLRs) possessing an N-terminal Toll-interleukin-1 receptor-like domain (TNLs). Here, immune functions of EDS1 complexes were comparatively analyzed in a mixed species approach relying on *Nicotiana benthamiana (Nb), Solanum lycopersicum (Sl)* and *Arabidopsis thaliana* (At). Genomes of most *Solanaceae* plants including *Nb* and *Sl* encode for two SAG101 isoforms, which engage into distinct complexes with EDS1. By a combination of genome editing and transient complementation, we show that one of these EDS1-SAG101 complexes, and not an EDS1-PAD4 complex as previously described in At, is necessary and sufficient for all tested TNL-mediated immune responses in *Nb.* Intriguingly, not this EDS1-SAG101 module, but mainly *Solanaceae* EDS1-PAD4 execute immune functions when transferred to At, and TNL functions are not restored in *Nb* mutant lines by expression of *At* EDS1 complexes. We conclude that EDS1 complexes do not represent a complete functional module, but co-evolve with additional factors, most likely protein interaction partners, for their function in TNL signaling networks of individual species. In agreement, we identify a large surface on SlEDS1 complexes required for immune activities, which may function in partner recruitment. We highlight important differences in TNL signaling networks between *At* and *Nb,* and genetic resources in the *Nb* system will be instrumental for future elucidation of EDS1 molecular functions.

## Introduction

Plants lack specialized mobile immune cells, but have evolved an elaborate innate immune system to defend against invading pathogens (Spoel and Dong, 2012; Jones et al., 2016). Cell surface-resident receptors termed pattern recognition receptors (PRRs) can detect pathogen-/ microbe-associated molecular patterns (PAMPs/MAMPs). MAMPs are often generic molecules common to both pathogenic and non-pathogenic microbes, such as chitin, peptidoglycan, lipopolysaccharides, or peptides derived from elongation factor EF-Tu or flagellin (Macho and Zipfel, 2014; Yu et al., 2017). MAMP perception and PRR activation induces PRR-triggered immunity (PTI; also referred to as MAMP/PAMP-triggered immunity, MTI/PTI), a multifaceted, low-level immune response efficient against a broad spectrum of non-adapted pathogens. However, host-adapted pathogens employ effector proteins, which are secreted directly into the host cell cytoplasm, to suppress or evade PTI (Macho and Zipfel, 2015; Büttner, 2016). As a second layer of the plant immune system, effectors can become recognized by plant *Resistance* proteins (R proteins) in resistant isolates of the host. Effector recognition induces a rapid and efficient immune response termed effector-triggered immunity (ETI) (Jones et al., 2016; Khan et al., 2016). The ETI response commonly culminates in programmed cell death at attempted infection sites, the hypersensitive response (HR), and HR-induction correlates in most cases with inhibition of microbial colonization (Cui et al., 2015; Büttner, 2016).

Most plant R proteins belong to the nucleotide-binding/ leucine-rich repeat (NLR)-type of immune receptors. The canonical NLR architecture consists of a C-terminal leucine-rich repeat (LRR) domain, a central nucleotide-binding (NB) domain and a variable N-terminal domain (Monteiro and Nishimura, 2018). Structurally similar NLR receptors are found in animal innate immunity, and can function by ligand-dependent oligomerization, followed by recruitment of signaling adapters via oligomeric N-terminal domain assemblies (Bentham et al., 2017). Plant NLRs are less well understood. However, in a general working model, the LRR domain often defines specificity of a respective NLR receptor, the central NB domain acts as an ATP-driven switch controlling the transition of the receptor from a resting to an active signaling state, and the N-terminal domain conveys downstream signaling (Maekawa et al., 2011a; Takken and Goverse, 2012; Bernoux et al., 2016). As an example, allelic variants of the Arabidopsis Resistance protein RPP1 recognize variants of the effector ATR1 from different isolates of the oomycete pathogen *Hyaloperonospora arabidopsidis* (*Hpa*), and effector recognition correlates with association of the cognate effectors with the immune receptor LRR domains (Krasileva et al., 2010; Steinbrenner et al., 2015). Also, the LRR domains of plant Resistance proteins, particularly of those occurring as highly divergent gene clusters within genomes, are often under positive selection, indicative of close co-evolution of effectors and immune receptors in a molecular arms race (Mondragon-Palomino et al., 2002; Jacob et al., 2013; Baggs et al., 2017). However, direct binding of effectors to the LRR domain is only one possible proxy for receptor activation. Resistance proteins may also e.g. monitor the integrity of a host protein in the ‘guard’ and ‘decoy’ models, or contain non-canonical domains now referred to as ‘integrated decoys’ functioning as an interaction platform to perceive effectors (Sarris et al., 2016; Cesari, 2018). Irrespective of the actual detection mode, effector recognition most likely alters intramolecular domain interactions, thereby shifting the receptor from a closed, ADP-bound to the ATP-bound active conformation (Bernoux et al., 2016; Jones et al., 2016). The N-terminal domains of plant NLRs are typically TIR (Toll-interleukin-1 receptor) or CC (coiled-coil) domains. Beyond analogy animal NLRs, major support for a function of these N-terminal domains of plant NLRs in downstream signaling is provided, on the one hand, by the observation that expression of TIR or CC domains from different NLRs is sufficient to induce HR-like cell death (Swiderski et al., 2009; Bernoux et al., 2011; Maekawa et al., 2011b). On the other hand, there is an apparent bifurcation of signaling pathways downstream of CC-and TIR-type NLRs: While many CC-type NLRs (CNLs) require NDR1 (Non-race specific Disease Resistance 1) for induction of immunity, all known responses mediated by TIR-type NLRs (TNLs) are dependent on EDS1 (Enhanced disease susceptibility 1; Century et al., 1997; Aarts et al., 1998; Wirthmueller et al., 2007).

*EDS1* was identified in Arabidopsis in a genetic screen for mutants impaired in resistance to the obligate biotrophic oomycete *Hpa,* and encodes for a protein with similarity to eukaryotic lipases in its N-terminus (Parker et al., 1996; Falk et al., 1999). EDS1 directly interacts with two sequence-related proteins, PAD4 (Phytoalexin deficient 4) and SAG101 (Senescence-associated gene 101) (Feys et al., 2001; Feys et al., 2005). All three proteins share the homology to eukaryotic lipases (α/β-hydrolases) in their N-termini, and furthermore contain a C-terminal EDS1-PAD4 (EP) domain. Indeed, presence of the unique EP domain together with an N-terminal lipase-like domain is the defining feature of the EDS1 family (Wagner et al., 2013). Critical residues of a catalytic triad (S-D-H including a characteristic GXSXG motif) are conserved in the lipase-like domain of EDS1 and PAD4 orthologs, but were not required for EDS1-PAD4 immune functions when tested, thus suggesting a non-catalytic mode-of-action (Wagner et al., 2013). A recently solved crystal structure of an EDS1-SAG101 heterodimer and a derived EDS1-PAD4 homology model showed that EDS1 engages into mutually exclusive heterodimers with PAD4 or SAG101 (Wagner et al., 2013), which differentially contribute to immunity in Arabidopsis: Loss of EDS1-PAD4 complexes (in *pad4* mutant plants) severely impairs immune signaling, while loss of EDS1-SAG101 (in *sag101* mutant plants) is largely compensated by presence of EDS1-PAD4 (Feys et al., 2005; Wagner et al., 2013). Complete loss of EDS1-based complexes (in *eds1* single or *pad4 sag101* double mutant plants) fully abolishes TNL-mediated resistance signaling. In agreement with genetic data, structure-guided mutations untethering EDS1 from PAD4 and SAG101 provided strong evidence for only heterodimeric assemblies executing immune functions (Wagner et al., 2013).

Deviant from the strict and undisputed requirement of EDS1 for TNL-mediated immune responses, EDS1 was reported to also contribute to resistance mediated by some CNLs and to basal resistance (Wiermer et al., 2005; Venugopal et al., 2009; Cui et al., 2017). To that end, for example *Pseudomonas syringae* (*Pst*) strain DC3000 bacteria grow to significantly higher titers in *eds1* mutant than in wild type lines (basal resistance). Similarly, Arabidopsis accession Col-0 (Col) is resistant to *Pst* bacteria translocating AvrRpt2, recognized by the CNL RPS2 (Bent et al., 1994; Mindrinos et al., 1994), but resistance is impaired in lines lacking EDS1 and the defense-associated hormone salicylic acid (SA; Venugopal et al., 2009; Vlot et al., 2009; Cui et al., 2017). However, *Pst* assumingly translocates up to 29 different effectors into host plants (Wei et al., 2015; Wei et al., 2018), and infection assays fail to detect weak recognition events in interactions termed ‘compatible’. Indeed, basal resistance is also impaired in lines lacking RAR1 or SGT1 (required for maturation and accumulation of NLRs (Kadota et al., 2010; Zhang et al., 2010)), supporting the occurrence of weak recognition in the *Pst* DC3000 – Col interaction. Thus, *eds1* basal resistance phenotypes might be explained by inactivation of the TNL sector and loss of one or several weak recognition events. Similarly, susceptibility of EDS1- and SA-deficient plants to otherwise ‘incompatible’ *Pst* AvrRpt2 bacteria can be interpreted as the combined loss of weak recognition (EDS1) and bolstering of immune responses via the SA sector without any direct contribution of EDS1 to CNL-mediated responses. This is supported by the observation that EDS1 and SA function additively, and not redundantly, in RPS2-mediated resistance, and by residual induction of cell death by AvrRpt2 even in absence of EDS1 and SA (Cui et al., 2017). Based on these assumptions, EDS1 complexes might not function beyond TNL-mediated resistance in microbial immunity. This is in conflict with conservation of EDS1-PAD4 in genomes of monocotyledonous plants and the eudicots *Aequilegia courulea* and *Mimulus guttatus,* which lost TNLs in the course of evolution (Collier et al., 2011; Jacob et al., 2013; Wagner et al., 2013). Therefore, it was proposed that EDS1-PAD4 form an ancient module regulating basal resistance, which has been co-opted for TNL-mediated immunity in eudicots (Feys et al., 2005; Rietz et al., 2011).

Similar to TIR domain signaling partners remaining elusive, also the molecular functions of EDS1 complexes in this process remain unresolved. Genetic screens so far failed to identify informative alleles, indicating genetic redundancy among signaling partners, lethality of respective alleles or a direct signaling mechanism. Notably, nuclear localization of EDS1 was shown to be necessary and also sufficient for several tested resistance responses at least in Arabidopsis (Garcia et al., 2010; Stuttmann et al., 2016). It is thus conceivable that EDS1 might directly control defense-related transcriptional outputs in a TNL-dependent manner, and EDS1 was reported to associate with a number of different TNLs (Bhattacharjee et al., 2011; Heidrich et al., 2011; Kim et al., 2012). However, association of EDS1 with chromatin was not reported, and the physiological relevance of EDS1-TNL interactions remains unclear (Sohn et al., 2012; Huh et al., 2017).

An available body of knowledge on the *EDS1* family stems almost exclusively from analyses in *Arabidopsis thaliana.* Here, we established *Nicotiana benthamiana* as a novel system for functional analysis of these genes, and their role in plant immunity. *N. benthamiana* lines defective in EDS1 or PAD4 and SAG101 were not altered in susceptibility to pathogenic *Xanthomonas* bacteria, which contradicts a direct role of these genes in basal immunity. Furthermore, we show that EDS1-SAG101, but not EDS1-PAD4, are necessary and sufficient for TNL-mediated resistance responses in *N. benthamiana.* On the basis of differential requirements for PAD4 and SAG101 in Arabidopsis and *N. benthamiana,* respectively, functional conservation of EDS1 complexes was probed by cross-species transfer. Intriguingly, *Solanaceae* EDS1-PAD4 (from tomato, *Sl)* fail to execute immune functions in *N. benthamiana,* but are fully functional in Arabidopsis. In contrast, SlEDS1-SlSAG101 function in *N. benthamiana,* but not in Arabidopsis, and *At*EDS1-*At*PAD4-AtSAG101 do not have any activity in *N. benthamiana.* We conclude that EDS1 complexes do not form a functional module themselves, but co-evolve with additional signaling partners, most likely protein interaction partners, in individual species. This is further supported by identification of several residues required for immune functions delineating a surface of the EDS1-SAG101 heterodimer, which extends from the N-terminal hydrolase to the C-terminal EP domains and might function in interaction partner recruitment. The newly established *N. benthamiana* system, in which TNL activation and signaling can be uncoupled from complex biotic interactions for rapid functional analyses, and also identified non-functional EDS1-SAG101 alleles, will prove seminal for future elucidation of EDS1 molecular functions.

## Results

### Duplication of SAG101 in *Solanaceae*

We previously generated *eds1* and *pad4* mutant lines in *N. benthamiana* (Ordon et al., 2017). In accordance with observations from the Arabidopsis system, plants lacking *NbEDS1* failed to restrict growth of *Xanthomonas campestris* pv. *vesicatoria* (strain 85-10, in the following Xcv) bacteria translocating the effector XopQ, which is recognized by the TNL Roq1 (Adlung et al., 2016; Schultink et al., 2017). However, *Xcv* growth was restricted in *Nbpad4* mutant plants as efficiently as in wild type plants (see later sections), suggesting lower importance of EDS1-PAD4 complexes in this system. This prompted us to further analyze the *EDS1* gene family in the genus *Solanaceae.* We searched *EDS1* family genes in the high quality genome of diploid tomato *(Solanum lycopersicum, Sl)* using Arabidopsis proteins as query (tBLASTn). Single copy genes for *EDS1* and *PAD4* (Adlung et al., 2016), and two different *SAG101* isoforms, which we termed *SAG101a* and *SAG101b,* were identified (Figure S1 and Table S1). Using the tomato proteins as query, further sequenced *Solanaceae* genomes were mined for *EDS1* family genes. In *N. benthamiana, PAD4* is encoded by a single gene, while a pseudogene and a functional copy were detected for *EDS1,* as previously described (Adlung et al., 2016). Similar to tomato and in accordance with allotetraploidy of *N. benthamiana,* two copies each of *SAG101a* and *SAG101b* were detected (Figure S1 and Table S1). Indeed, *SAG101a* and *SAG101b* isoforms were detected in all analyzed *Solanaceae* genomes (Table S1) except pepper *(Capsicum annuum).* Phylogenetic clustering of respective proteins from *Solanaceae,* Arabidopsis and control species revealed that EDS1 and PAD4 homologs grouped together, respectively, while SAG101a and SAG101b formed two distinct groups within the SAG101 branch (Figure 1a). The single copy SAG101 from pepper grouped within the SAG101b branch. SAG101 isoforms from outside the *Solanaceae (Coffee canephora* (Cc) and Arabidopsis) were more closely related to, but did not directly cluster with, SAG101b (Figure 1a). SAG101 homologs were not detected in species lacking TNL receptors (here *Musa accuminata (Ma)* and *Mimulus guttatus (Mg)),* as previously described (Wagner et al., 2013).

**Figure 1:**
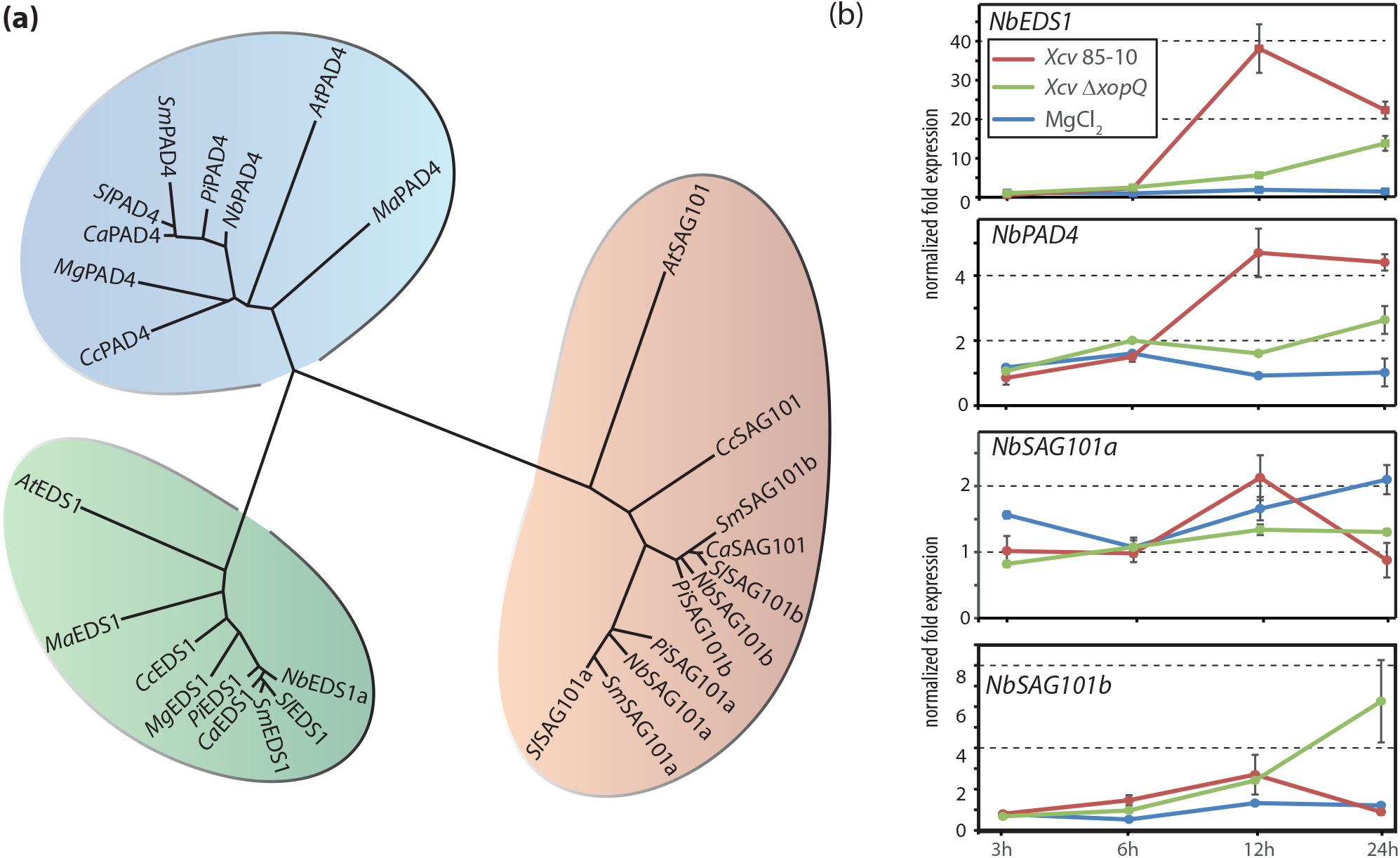
Occurrence and expression of *EDS1* family genes in *Solanaceae*. a) Phylogenetic clustering of putative EDS1 family proteins from *Solanaceae* and control species in radial representation. EDS1, PAD4 and SAG101 branches are marked in green, blue and light brown, respectively. *At – Arabidopsis thaliana; Ma – Musa accuminata; Cc – Coffee canephora; Mg – Mimulus guttatus; Pi – Petunia infìata; Ca – Capsicum annuum; Sm – Solanum melongena; Sl – Solanum lycopersicum; Nb – Nicotiana benthamiana.* b) Expression of *EDS1* family genes in *Nicotiana benthamiana.* Plants were challenged with virulent *(XcvΔxopQ)* or avirulent (Xcv 85-10) *Xanthomonas campestris* pv. *vesicatoria* bacteria, or mock (MgCl_2_)-treated. RNA was extracted at indicated time points, and expression of *EDS1* family genes measured by quantitative RT-PCR. Displayed data originates from normalization to *Protein Phosphatase 2A (PP2A)* expression. Data points represent means of four biological replicates with standard error shown.

A public RNAseq dataset from plants treated with different bacterial isolates or the MAMP flgII-28 (Rosli et al., 2013) was assessed using the TomExpress platform (Zouine et al., 2017) to analyze expression of tomato *EDS1* family genes (Figure S2). Transcription of all four genes was detected in mock-treated plants, confirming general expression of both *SAG101* isoforms. Treatment with *Pseudomonas fluorescens* (*Pfo* 55) and *Pseudomonas putida* bacteria, inducing robust MTI responses in *Solanaceae* (Chakravarthy et al., 2010; Rosli et al., 2013), moderately induced expression of *EDS1* family genes (6 hours post treatment). We further examined the expression of *EDS1* family genes in *N. benthamiana.* The *SAG101a2/b2* isoforms were omitted from analysis, as we assumed them to be pseudogenes due to altered intron-exon structure of the respective gene models (Figure S1). Plants were challenged with avirulent *(Xcv)* or virulent (Xcv *ΔxopQ) Xanthomonas* bacteria, and gene expression analyzed by quantitative RT-PCR in a time course experiment (Figure 1b). Expression of *NbEDS1* was strongly upregulated in response to avirulent *Xcv* (recognized by presence of XopQ), and to lesser extent by the virulent *ΔxopQ* mutant strain (Figure 1b). Expression peaked 12h post infection, and subsequently declined. A similar trend was observed for *NbPAD4,* although its overall induction (~35-fold for *NbEDS1,* 5-fold for *NbPAD4;* Figure 1b) was less pronounced. *NbSAG101a* and *NbSAG101b* were also expressed, but were not strongly regulated under infection conditions. Overall, these data suggest that two different SAG101 isoforms are expressed at least in tomato and *N. benthamiana,* which may engage into heterocomplexes with EDS1 and contribute to immune signaling.

### Localization and complex formation of tomato EDS1 family proteins

We aimed to analyze EDS1 functions in *N. benthamiana* as a model system. However, *EDS1* family genes were first cloned from tomato for its high quality genome and reduced complexity of the gene family. SlEDS1 was fused to mCherry and SlPAD4, SlSAG101a and SlSAG101b to mEGFP, respectively, in 35S promoter-controlled expression constructs. Fusions proteins were, alone or in combination, transiently expressed in *N. benthamiana,* and subcellular localization was analyzed by live cell imaging (Figures 2a, S3). As previously described for Arabidopsis orthologues (Feys et al., 2005; Garcia et al., 2010), *Sl*EDS1 and *Sl*PAD4 localized to both the cytoplasm and the nucleus. Similarly in analogy to the Arabidopsis system, SlSAG101a located exclusively to the nucleus. In co-expression with SlEDS1, SlSAG101a re-localized SlEDS1 to the nucleus (compare Figures 2a and S3a). In contrast, SlSAG101b was nucleo-cytoplasmically distributed both alone and in combination with SlEDS1. All proteins were detected on immunoblots, and the GFP-tagged SlPAD4, SlSAG101a and SlSAG101b appeared to be stabilized by co-expression of SlEDS1 (Figure S2b). Förster resonance energy transfer and acceptor photobleaching (FRET-APB) was used to assess formation of SlEDS1-based complexes in living cells. mCherry and mEGFP-tagged proteins were expressed from a single T-DNA for reduced variation in co-expression rates (Hecker et al., 2015). Robust FRET was detected upon co-expression of SlEDS1-mCherry and either SlPAD4, SlSAG101a or SlSAG101b, but not SlEDS1, fused with mEGFP (Figure S3c). Complex formation was further analyzed by protein co-purification (Figure 2b). In line with results obtained by FRET-APB, SlEDS1 (fused to a 6xHA tag) co-purified with Strep-tagged SlPAD4, SlSAG101a and SlSAG101b, but not SlEDS1. We conclude that SlEDS1 engages into three different heterocomplexes containing SlPAD4, SlSAG101a or SlSAG101b, but does not form homodimers. SlEDS1 family proteins are highly similar to those of *N. benthamiana* (79/86 (EDS1), 77/85 (PAD4), 81/87 (SAG101a) and 72/79 (SAG101b) % identity/similarity). Thus, at least three different EDS1-based heterocomplexes are most likely also expressed in *N. benthamiana*, and one or several of these are expected to function in TNL immune signaling.

**Figure 2:**
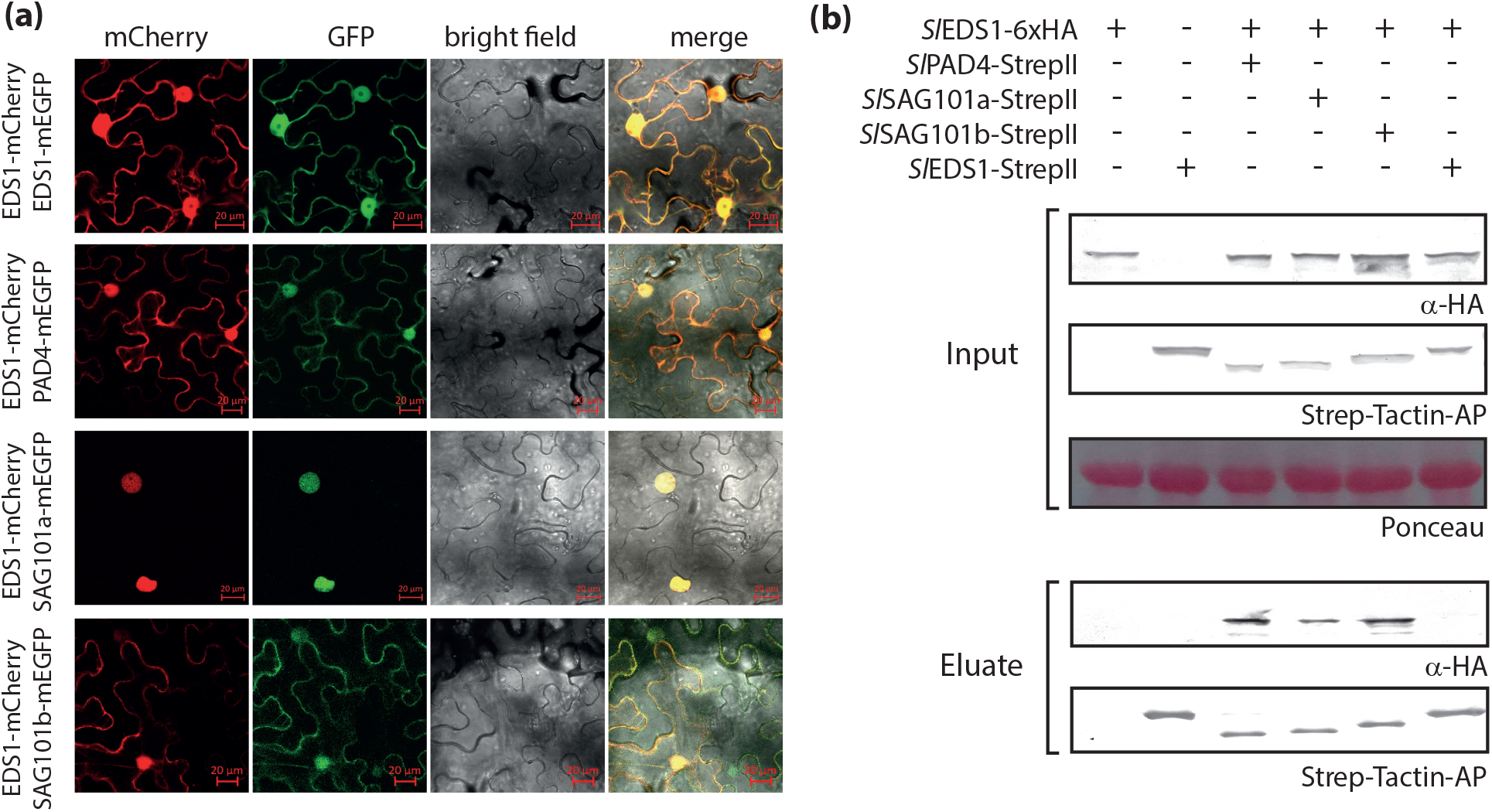
Complex formation and localization of tomato EDS1 family proteins. a) Protein localization in living cells detected by confocal laser scanning microscopy. Indicated proteins (from tomato) were transiently co-expressed in *N. benthamiana* leaf tissues by Agroinfiltration, and protein localization was analyzed 3 dpi. Localization of single proteins and integrity of fìuorophore fusions is shown in Figure S3. b) Formation of complexes by tomato EDS1 proteins. Indicated proteins were transiently (co-) expressed in *N. benthamiana* by Agroinfiltration. At 3 dpi, extracts were used for Strepll-purification, and total extracts and eluates analyzed by immunoblotting. Ponceau-staining is shown as loading control.

### Identification of EDS1 complexes functioning in XopQ recognition

Recognition of XopQ delivered as an AvrRpt2 fusion by *Pseudomonas fluorescens (Pfo XopQ)* induces a strong HR response in *N. benthamiana* (Figure 3a; Gantner et al., 2018). The XopQ-induced HR was abolished on *eds1* plants, but not affected by mutation of *PAD4* (Figure 3a). We assumed that one or several of the NbSAG101 isoforms (Figures 1a, S1) might function redundantly with NbPAD4 in TNL-mediated resistance. To identify respective *NbSAG101* isoforms, the previously generated *pad4-1* mutant line (Ordon et al., 2017) was transformed with a genome editing construct for disruption of all four *SAG101a* and *SAG101b* copies (Figure S4). Primary transformants (T_0_) were directly screened for recognition of XopQ by inoculation with *Pfo XopQ.* One plant unable to initiate HR in response to XopQ, and thus phenocopying the *Nbeds1* line, was identified. Sequencing revealed that *SAG101a2* and *SAG101b2* did not contain any mutations in this line, and can be dismissed for immune functions. Since we assume these copies to represent pseudogenes, only *SAG101a1/b1* will be considered in the following, and will be referred to as *SAG101a and SAG101b* for simplicity. The primary line non-responsive to XopQ was homozygous for a *sag101a-1* mutation, and heterozygous for two different *sag101b* alleles (see Figure S4 for details). A line void of the genome editing transgene and homozygous for disruptive alleles at *SAG101a/b* (thus a *pad4-1 sag101a-1 sag101b-1* triple mutant; *pss)* was selected. A further double mutant line containing the *pad4-1* and *sag101b-1* mutant alleles (pSs) was isolated from a cross *(pss* x wild type). We failed to isolate a *sag101b-1* single mutant line from the same cross. Allele identifiers will be omitted for *N. benthamiana* lines in the following.

**Figure 3:**
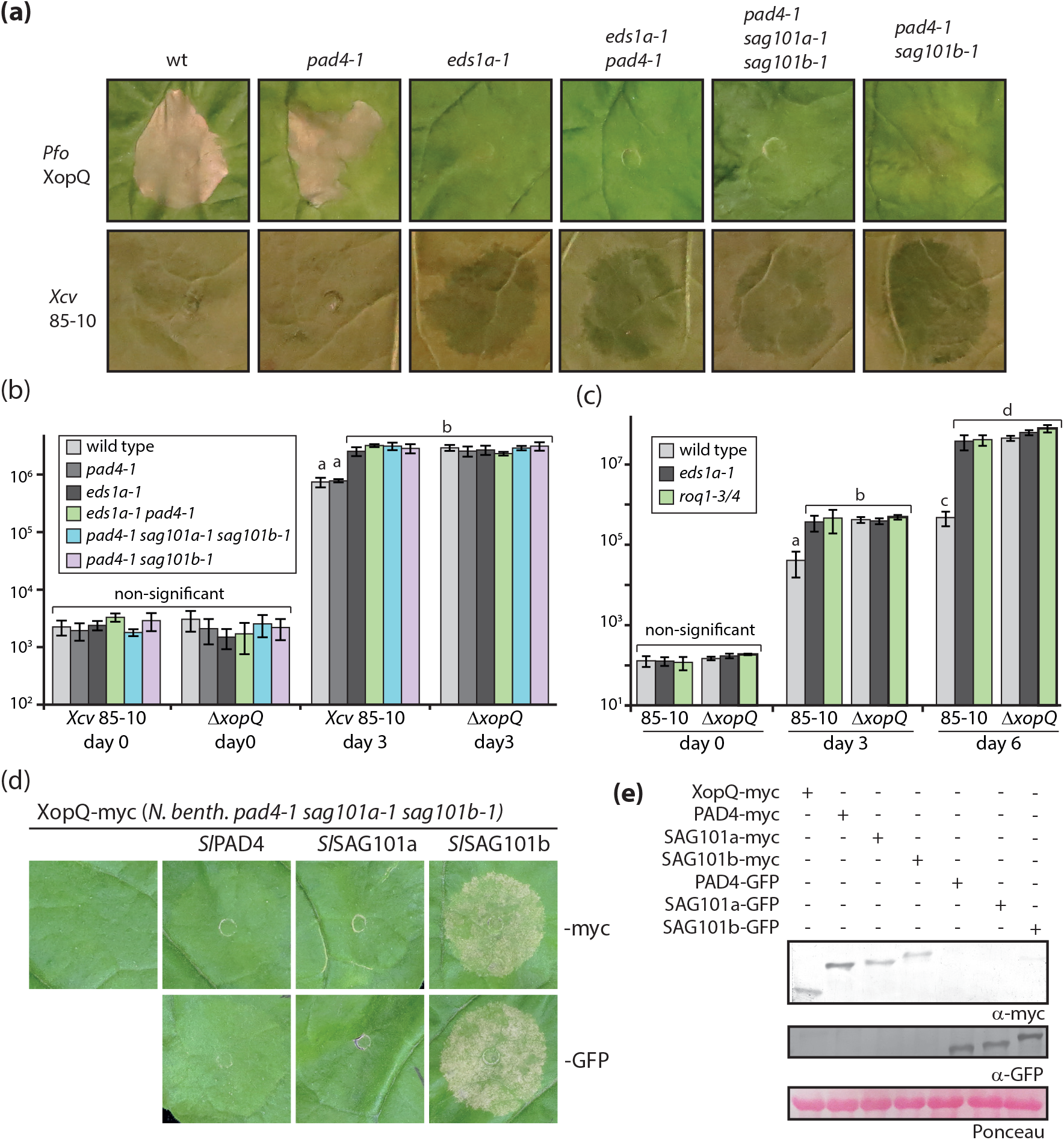
Immune responses of *Nicotiana benthamiana* mutant lines deficient in *EDS1* family genes or *Roq1*. a) Recognition of XopQ in different mutant lines. Indicated *N. benthamiana* lines were challenged with XopQ-translocating *Pseudomonas fluorescence* bacteria (upper panel; infiltrated at OD_600_ = 0.2) or *Xcv* strain 85-10 bacteria (lower panel; infiltrated at OD_600_ = 0.4). Phenotypes were documented at 4 dpi. b) Bacterial growth of *Xcv* bacteria on mutant lines. Indicated lines were infected with *Xcv* strain 85-10 or a corresponding mutant strain lacking *XopQ (ΔxopQ).* Means and standard deviation of four biological replicates are shown. Letters indicate statistically significant differences as determined by one way ANOVA and Fisher LSD post hoc test (p < 0.01). c) Bacterial growth in *eds1* and *roq1* mutant lines. As in b), but means and standard deviation of eight biological replicates is shown for day 3 and day 6. The *roq1* mutant line was a T_1_ line segregating for two disruptive alleles at the *Roq1* locus (see Figure S4f for details). d) Reconstitution of XopQ detection in the *pss* triple mutant line. By Agroinfiltration, XopQ was expressed alone or in combination with either PAD4, SAG101a or SAG101b (from tomato and fused to a Twin-Strep and 4 x c-myc tag or GFP). Phenotypes were documented 5 dpi. e) Immunodetection of fusion proteins used in d). Individual fusion proteins were expressed in *pss* mutant plants, and detected in protein extracts 3 dpi. Ponceau-staining of the membrane is shown as loading control.

Mutant lines deficient in *EDS1* family genes were tested alongside control plants for recognition of XopQ by inoculation with *Pfo XopQ* and *Xcv* bacteria (Figure 3a). Wild type and *pad4* plants developed an HR, visible by strong cell death upon *Pfo XopQ* challenge and absence of disease symptoms in case of *Xcv* infection. The remaining mutant lines *(eds1 pad4 (ep), pss, pSs)* were unable to initiate HR and phenotypically indistinguishable from *eds1* mutant plants (Figure 3a). *In planta* bacterial titers of *Xcv* and *Xcv ΔxopQ* bacteria (eliciting Roq1- and EDS1-dependent immunity via XopQ or not, respectively) were determined for quantitative analysis of immune responses in different *N. benthamiana* mutant lines (Figure 3b). The growth of *Xcv,* but not *ΔxopQ* bacteria, was restricted on wild type and *pad4* plants. We did not detect differences for *Xcv* replication between *eds1* plants and any of the *pad4 sag101* mutant lines (Figure 3b). Conclusively, the *SAG101a* isoform present in the *pad4 sag101b (pSs)* double mutant does not contribute to immunity, and loss of *PAD4* and *SAG101b* phenocopies *eds1* mutant lines. Notably, none of the mutant lines showed an enhanced susceptibility / basal resistance phenotype: Bacterial titers of *ΔxopQ* (on any plant genotype) and *Xcv* on *eds1* or *pad4 sag101* mutant plants were identical. We further compared bacterial growth in *eds1* and *roq1* mutant plants (see Figure S4 for details on *roq1* mutant) in a time course extended up to six days (Figure 3c). Identical bacterial titers (Xcv) were observed in *eds1* and *roq1* plants, and were not different from bacterial growth of *ΔxopQ* bacteria. This confirms that plants deficient in EDS1 complexes do not have any basal resistance phenotype in *N. benthamiana,* and that XopQ is indeed the only *Xcv* effector inducing (via Roq1) *EDS1*-dependent defenses (Adlung et al., 2016).

Full susceptibility of *pad4 sag101b* mutant lines to *Xcv* suggests that EDS1-PAD4 and EDS1-SAG101b complexes might function redundantly in TNL-mediated immune signaling. Alternatively, only EDS1-SAG101b might have immune functions in *N. benthamiana.* Since we had failed to isolate *sag101b* single mutants, transient expression of XopQ and tomato SlPAD4 or SlSAG101 proteins in the *pss* background was used to discriminate between these scenarios (Figure 3d). Only co-expression of SlSAG101b, but not SlPAD4 or SlSAG101a, together with XopQ restored HR induction. All proteins were detected on immunoblots using two different epitope tags (Figure 3e). The same transient complementation assay was performed with untagged proteins with identical results. We also compared NbEDS1 and NbSAG101b with respective tomato orthologues for restoration of XopQ recognition (Figure S5). No qualitative differences were detected (Figure S5a,b), and SlPAD4, SlSAG101a and SlSAG101b engaged into complexes with NbEDS1 (Figure S5c). We concluded that tomato EDS1 family proteins can functionally replace *N. benthamiana* orthologues, and decided to further use tomato proteins for functional characterization. Taking together that *i) pad4* mutant *N. benthamiana* lines are not impaired in XopQ-mediated resistance (Figures 3a,b) and that *ii)* only expression of SAG101b (Figure 3d) can restore XopQ recognition in the *pss* background, these results suggest that an EDS1-SAG101b complex is necessary and sufficient for resistance signaling downstream of XopQ in *N. benthamiana.* Despite its marked upregulation under infection conditions (Figure 1b), *PAD4* appears not to contribute to immune signaling in *N. benthamiana,* or at least is unable to function in absence of SAG101b.

### EDS1-SAG101b functions in diverse TNL-mediated responses

The observation that EDS1 and SAG101b are required for XopQ-induced resistance responses was surprising, as major resistance signaling functions reside in EDS1-PAD4 in Arabidopsis (Feys et al., 2005; Wagner et al., 2013; Cui et al., 2017; Cui et al., 2018). We tested additional inducers of assumedly EDS1-dependent defense responses in our set of mutant lines to analyze whether EDS1-SAG101b are generally required for immune signaling in *N. benthamiana*, or whether this might be specific for the TNL Roq1 recognizing XopQ (Schultink et al., 2017). Expression of a TIR domain fragment of the TNL DM2h (Stuttmann et al., 2016), a TIR fragment of RPS4 (Swiderski et al., 2009; Williams et al., 2014) and co-expression of the TMV Helicase protein p50 together with the tobacco TNL receptor N (Burch-Smith et al., 2007) induced HR-like cell death on wild type, but not *eds1* or *pss* mutant plants (Figure 4a). As with XopQ, cell death formation could be restored in *pss* plants by co-expression of SlSAG101b, but not SlSAG101a or SlPAD4. These results suggest that EDS1-SAG101b is generally required for TNL-induced defenses in *N. benthamiana,* while PAD4 and SAG101a cannot functionally replace SAG101b even when expressed to high levels in transient assays.

**Figure 4:**
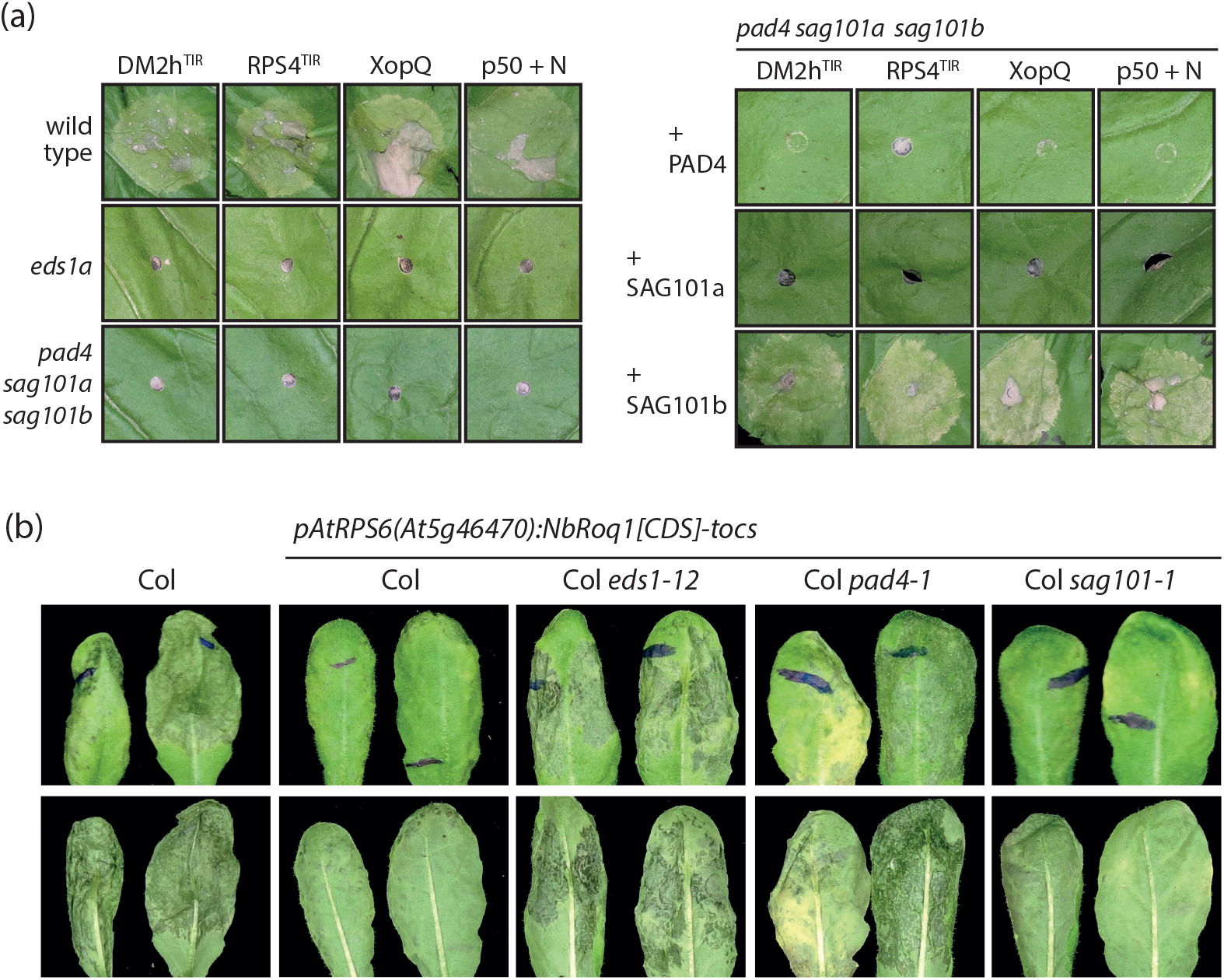
Genetic dependencies of TNL-type immune receptors in *N. benthamiana* and Arabidopsis. a) EDS1-dependent cell death induction requires SAG101b in *N. benthamiana.* Inducers of presumably EDS1-dependent cell death (DM2h^TIR^ – DM2h(_1-279_); RPS4^TIR^ – RPS4(_1-234_)_E111K (Swiderski et al., 2009); XopQ – XopQ-myc; p50 + N – p50-Cerulean + N-Citrine (Burch-Smith et al., 2007)) were expressed in different *N. benthamiana* lines as indicated (left panel), or co-expressed with PAD4, SAG101a or SAG101b (from tomato and fused to a 4xmyc-TwinStrep tag) in the *pss* mutant line (right panel). Phenotypes were documented 5 dpi. b) Functionality and genetic dependency of Roq1 in *Arabidopsis thaliana.* A T-DNA construct coding for Roq1 under control of an *RPS6* promoter fragment and an *ocs* terminator was transformed into the indicated Arabidopsis lines. Four week-old control and T_1_ plants were infected with *Pst* DC3000 bacteria (syringe infiltration, OD_600_ = 0.001). Symptom development was documented 3 dpi. At least eight independent T_1_ plants were tested for each genotype per replicate, and the experiment was repeated once with similar results.

In a complementary approach, the *Roq1* gene was transferred into Arabidopsis wild type and *eds1, sag101* and *pad4* mutant lines. *Roq1* was expressed under control of the promoter of Arabidopsis *RPS6* and the *ocs* terminator *(octopine synthase, Agrobacterium tumefaciens). RPS6* encodes for a TNL receptor recognizing HopA1 from *Pseudomonas syringae* pv. *syringae* strain 61 (Kim et al., 2009), and its promoter was chosen for *Roq1* expression to potentially avoid dominant negative effects often arising from overexpression of immune receptors (e.g. Wirthmueller et al., 2007). T_1_ transgenic seeds from transformation of the *Roq1* expression constructs into Columbia (Col-0) wild type and *eds1-12, pad4-1* and *sag101-1* mutant lines were selected by FAST seed coat fluorescence (Shimada et al., 2010), respective plants grown in soil and directly infected with *P. syringae* pv. tomato *(Pst)* strain DC3000. DC3000 contains the effector HopQ1 homologous to XopQ, which is also recognized by Roq1 (Adlung et al., 2016; Schultink et al., 2017; Zembek et al., 2018). We therefore expected to generate resistance to *Pst* DC3000, which is highly virulent on Col-0, if Roq1 can function in Arabidopsis. Indeed, severe tissue collapse was observed in Col-0 plants infected with DC3000 at 3dpi, while plants containing the *Roq1* transgene were mostly asymptomatic (Figure 4b). As a TNL receptor, we expected Roq1 to function in an EDS1-dependent manner in Arabidopsis. In agreement, tissue collapse similar to that of Col-0 plants was observed in *eds1-12 pRPS6:Roq1* plants. The *pad4-1* transgenics containing the *pRPS6:Roq1* transgene were similar to *eds1-12* transgenics and Col-0, while transgenic lines in the *sag101* background were as resistant as wild type plants expressing Roq1 (Figure 4b). Similar results were obtained when Roq1 was expressed under the control of a *Ubiquitin 10* promoter fragment. Roq1 is thus functionally dependent on EDS1-SAG101b in *N. benthamiana,* but requires EDS1-PAD4 to mediate resistance in Arabidopsis. We conclude that not the TNL receptors, but differences within the EDS1 protein family of respective plant species, decide which EDS1 heterocomplexes function in TNL signaling.

### EDS1 complexes are not sufficient for TNL signaling, but additional factors divergent between individual species are required

Considering that either EDS1-PAD4 or EDS1-SAG101b are sufficient for immune signaling in Arabidopsis and *N. benthamiana,* respectively, these complexes might have identical functions, albeit different evolutionary origins. Alternatively, functional recruitment to immune signaling might occur by different mechanisms in these species. We sought to analyze these aspects by transferring *EDS1* family genes from Arabidopsis to *N. benthamiana,* and *vice versa.* We first attempted to restore XopQ-induced cell death in *eds1* or *pss* mutant *N. benthamiana* plants. Arabidopsis *EDS1* family genes were expressed, in different combinations and with or without an epitope tag, from a single T-DNA, and XopQ was co-expressed (Figure S6a). As controls, SlEDS1-HA and SlSAG101b-myc were co-expressed with XopQ. Arabidopsis and tomato proteins were detected on immunoblots, and orthologues from different species were expressed to similar levels (Figure S6b). However, XopQ-induced cell death was restored by co-expression of SlEDS1 and SlSAG101b (in *eds1* and *pss* mutant plants, respectively), but not by co-expression of the Arabidopsis EDS1 family proteins, in any given combination (Figure 5a). Thus, Arabidopsis EDS1 complexes cannot function in Roq1 signaling in *N. benthamiana.* Reciprocally, the Arabidopsis *eds1-2 pad4-1* double mutant line was transformed with constructs encoding for EDS1 and PAD4 from either Arabidopsis or tomato and under control of the corresponding native promoter elements from Arabidopsis (Figure S6c). For each transformation, several independent T_2_ populations were tested for complementation of the *eds1-2 pad4-1* immunity defects by infection *with Hyaloperonospora arabidopsidis (Hpa)* isolate Cala2. Isolate Cala2 is recognized via the TNL RPP2 in Col-0 (Sinapidou et al., 2004), and is highly virulent on *eds1 pad4* deficient plants (Figure 5b). Transformants expressing EDS1-PAD4 from tomato were as resistant to *Hpa* Cala2 as transformants expressing the Arabidopsis homologs, and indistinguishable from wild type Col-0 (Figure 5b). In simultaneously generated transgenics expressing epitope-tagged variants, EDS1 and PAD4 from Arabidopsis and tomato accumulated to similar levels as assessed by immunodetection (Figure S6d). An additional set of transgenic plants was generated in the *eds1-2 pad4-1 sag101-1* triple mutant background that expressed SlEDS1 and different combinations of SlPAD4 and/or SlSAG101 isoforms (Figure S6e). Lines expressing SlEDS1 together with SlPAD4 and SlSAG101 isoforms were resistant to *Hpa* isolate Cala2, although hypersensitive response-associated cell death appeared less confined than in Col-0 or control lines expressing Arabidopsis EDS1-PAD4 (Figure S6e). In contrast, transgenics expressing SlEDS1 and a SlSAG101 isoform, but not SlPAD4, were susceptible. Again, all proteins were detected in simultaneously generated transgenics expressing epitope-tagged variants (Figure S6f). Hence, SlEDS1-SlSAG101b are sufficient for all tested immune responses in *N. benthamiana,* but fail to function in Arabidopsis. In contrast, SlEDS1-SlPAD4 can function in RPP2-mediated resistance in Arabidopsis, but not in any tested TNL-mediated response in *N. benthamiana.* Thus, proteins of the PAD4 phylogenetic clade appear to function, together with EDS1, in TNL signaling in Arabidopsis, and these functions are executed by EDS1-SAG101 in *N. benthamiana.* Taken together with the observation that *At*EDS1-*At*SAG101-AtPAD4 are not functional in *N. benthamiana,* we conclude that EDS1 complexes do not form a functional module in TNL signaling by themselves. We propose that additional factors, divergent between species as a result of co-evolution with EDS1 complexes, are required for these heterocomplexes to function in immunity.

**Figure 5:**
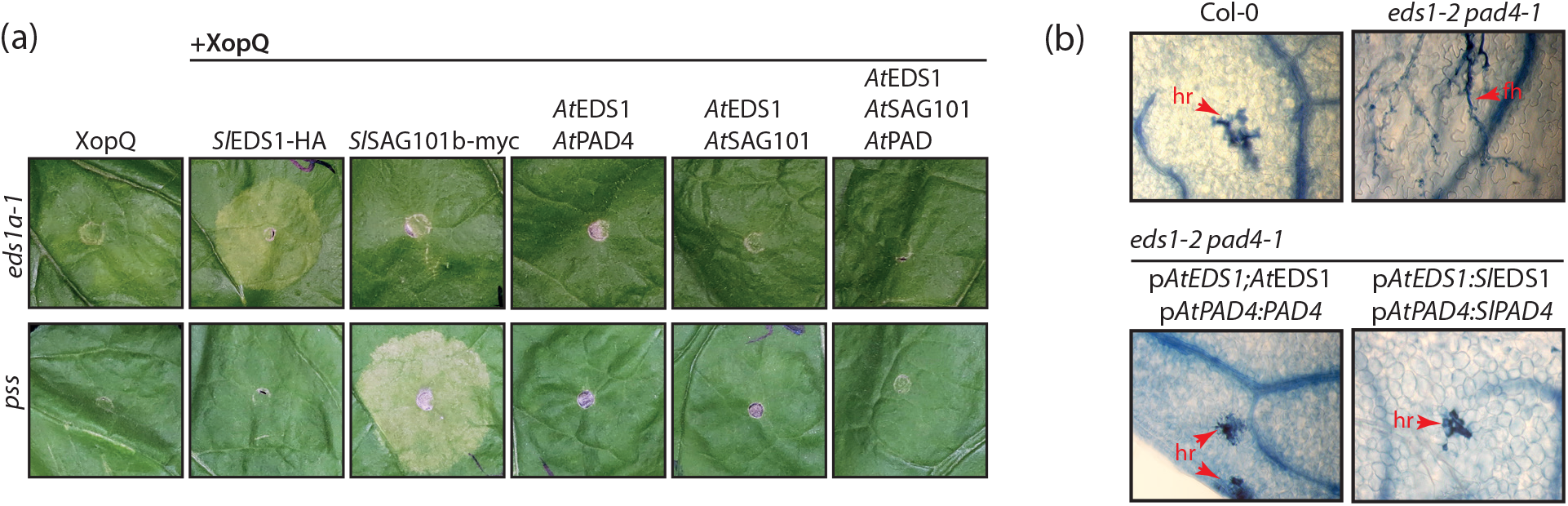
Cross-species transfer of *EDS1* family genes. a) Arabidopsis EDS1-PAD4-SAG101 proteins cannot functionally replace EDS1-SAG101b in *N. benthamiana.* Indicated proteins were expressed (by Agroinfiltration) either in *eds1* or *pss* mutant lines, and phenotypes were documented 7 dpi. Arabidopsis proteins were expressed with or without an epitope tag, and images originate from untagged proteins (see Figure S5 for details on T-DNA constructs and protein detection). b) Tomato EDS1-PAD4 can function in TNL-signaling in Arabidopsis. Col *eds1-2 pad4-1* double mutant was transformed with constructs for expression of EDS1 and PAD4, either from Arabidopsis or tomato and with or without an epitope tag (see Figure S5c for details) and under control of Arabidopsis promoter fragments. Segregating T_2_ populations were selected with BASTA, and three week-old plants infected with *Hyaloperonospora arabidopsidis* isolate Cala2. True leaves were used for Trypan Blue staining 7 dpi. At least four independent T_2_ populations were tested for each construct with similar results. Lines expressing untagged proteins were used for infection assays. Lines expressing epitope-tagged proteins were used for immunodetection (Figure S5d). hr – hypersensitive response; fh – free hyphae.

### Use of the *N. benthamiana* system for rapid functional analyses

One motive for genetic dissection of the *EDS1* gene family in *N. benthamiana* was the establishment of a novel experimental system allowing rapid functional analysis of these signaling components. Previous experiments had shown that XopQ-induced cell death can be restored in the *eds1* and *pss* mutant lines by *Agrobacterium*-mediated, transient co-expression of EDS1 or SAG101b, respectively (e.g. Figure 4a; Adlung et al., 2016; Qi et al., 2018). The HR-like cell death provoked by XopQ expression is relatively mild on wild type *N. benthamiana* plants and further delayed and dampened in transient complementation assays, but was highly reproducible under our conditions.

A crystal structure of the Arabidopsis EDS1-SAG101 complex and an experimentally validated homology model of the EDS1-PAD4 complex were previously reported (Wagner et al., 2013). Since Arabidopsis EDS1 complexes were not functional in *N. benthamiana*, homology models of EDS1-based heterodimers from tomato were generated. A structure similar to that of Arabidopsis EDS1-SAG101 was predicted for the functional tomato EDS1-SAG101b complex, and most surface-exposed, conserved residues mapped to the heterocomplex interface (Figures 6a, b). To validate the structural models and also the *N. benthamiana* system for functional analyses, we decided to first disrupt EDS1-SAG101b complex formation by mutagenesis of key amino acids within the N-terminal interaction interface (Wagner et al., 2013). The N-terminal interface is formed mainly by hydrophobic interactions between a protruding helix of EDS1 accommodated in a corresponding pocket on SAG101b (Figure 6a). Residues within the EDS1 helix were sequentially mutated: T264F and I268E (TI), followed by V265E (TIV), V269E (TIVV) and L261E (TIVVL). All variants accumulated to comparable levels *in planta,* and TIV or higher order mutants did not co-purify to detectable amounts with StrepII-tagged SAG101b *ex planta* (Figure 6c). When tested by yeast two hybrid (Y2H), interaction of the EDS1 variants with SAG101b and also PAD4 and SAG101a exhibited a gradual decline, and was still detectable for the TIVV quadruple mutant variant (Figure S7). In accordance with complex formation being progressively impaired, SlEDS1 variants also lost their activity for restoring XopQ-induced cell death when co-expressed in *eds1* mutant *N. benthamiana* plants, and only the quintuple TIVVL variant was completely non-functional (Figure 6d). Similarly, mutations were serially introduced into SAG101b: F17S and L22S (FL), L13S (FLL), L16S (FLLL) and L18S (FLLLL). In co-purification assays, interaction to EDS1 was detectable only for wild type SAG101b, and protein accumulation of SAG101 variants was only mildly affected (Figure 6e). SAG101b variants were tested for functionality by appearance of HR-like cell death upon co-expression with XopQ in *pss* mutant plants (Figure 6f). Cell death was reduced for SAG101b-FLL, and abolished for the quadruple and quintuple mutant variants. These data suggest that heterocomplex formation is required for immune functions of EDS1 and SAG101b, and thus fully confirm previous findings obtained in the Arabidopsis system (Wagner et al., 2013).

**Figure 6:**
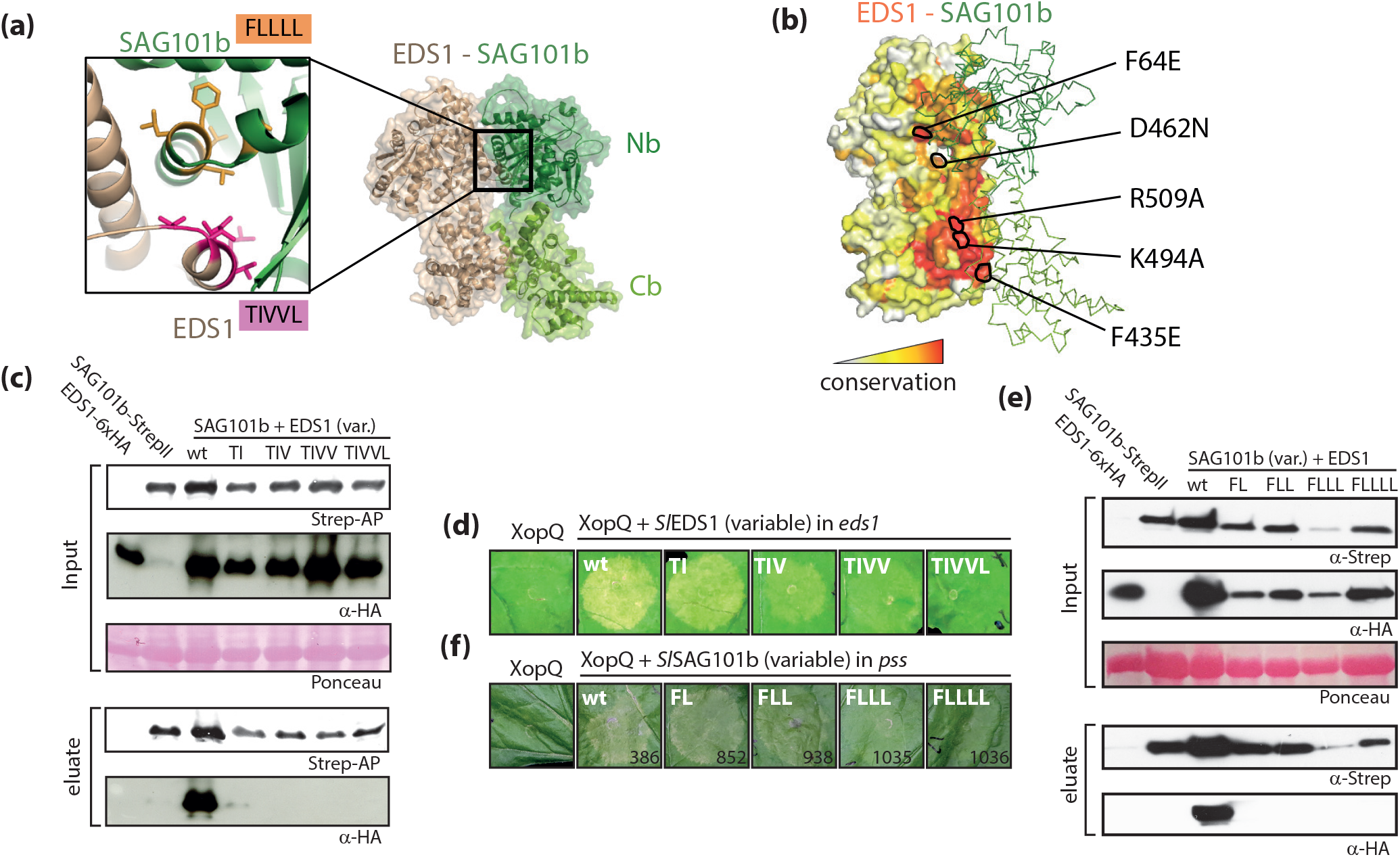
EDS1-SAG101b heterocomplexes are the functional modules in *N. benthamiana*TNL-signaling. a) Homology model of the tomato EDS1-SAG101b complex used for transient reconstitution of TNL-signaling in mutant *N. benthamiana* tissues. The inset shows the symmetrically arranged helices of EDS1 and SAG101 forming the N-terminal interaction interface. Amino acids targeted by mutagenesis are shown as sticks and are highlighted in pink (EDS1) and orange (SAG101), respectively. b) Conservation of surface-exposed amino acids in S/EDS1. S/SAG101 is shown in ribbon presentation (green). EDS1 residues functionally interrogated by mutagenesis are marked. c) Interaction of EDS1 variants with SAG101b. Indicated proteins were (co-) expressed in *N. benthamiana* by Agroinfiltration, and tissues used for StreplI-purification at 3 dpi. d) Functionality of EDS1 variants affected in heterocomplex formation. Indicated variants (as in d) with C-terminal 6xHA) were co-expressed with XopQ-myc in *eds1* mutant plants, and plant reactions documented 7 dpi. e) Interaction of SAG101 variants with EDS1. As in d), but SAG101-StrepII variants were co-expressed with EDS1. f) Functionality of SAG101 variants affected in heterocomplex formation. SAG101-StrepII variants were co-expressed with XopQ-myc in *pss* mutant plants, and plant reactions documented 7 dpi.

Having validated the transient *N. benthamiana* complementation assay for analysis of EDS1-SAG101b immune functions, we set out to identify additional functionally relevant features of these complexes (see Table 2 for a summary of tested variants). We first focused on several positively charged residues lining an assumed cavity on the heterodimer surface (Wagner et al., 2013) and recently reported as required for immune signaling in Arabidopsis (Bhandari et al., 2018). The residues within SlEDS1 (R509, K494) homologous to those reported in Arabidopsis (R493, K478) were targeted by mutagenesis, and respective variants tested for functionality (Figure S8). All variants restored XopQ-induced HR-like cell death as efficiently as wild type SlEDS1. We propose that functional relevance of the positively charged residues might be masked by over-expression in the *N. benthamiana* system, or might not be conserved across the different systems. We further introduced F64E, F435E and D462N exchanges into SlEDS1 (Figure 6b). F64 is a single, conserved residue exposed on the N-terminal lipase-like domain of EDS1 and framing the assumed cavity. F435 is fully buried by the association with SAG101 C-terminal domain and can used as a probe to test the importance of the interaction between the C-terminal domains. D462 connects the N and C-terminal domains together, and might mediate crosstalk at the domain interface. Only the D462N variant was mildly affected in protein accumulation, and all variants retained interaction with SAG101b as tested by co-purification (Figure 7a). When co-expressed with XopQ in *eds1* plants, immune activities were reduced for F64E and D462N variants, and fully abolished for F435E. We further tested F435D and F435A variants. While F435D also failed to restore immune capacity in *eds1* plants, F435A was fully functional (Figure S8). We assume that disruption of the apolar patch at the C-terminal interface and introduction of a charged residue in F435E/D dislocates the EP domains within the EDS1-SAG101b heterocomplex against each other, without disturbing overall complex assembly mainly driven by the N terminal interaction surface. This supports a function of the heterodimeric EP domain surface in TNL signaling. We constructed chimeric proteins from non-functional SAG101a and functional SAG101b to further analyze this aspect (Figure 7c). Although differences between SAG101a/b isoforms remain unclear, we hypothesized that, if the EP domain surface is crucial for immune functions, only the chimeric protein carrying the C-terminus of SAG101b might be functional, while both chimera should be able to engage into heterocomplexes with EDS1. SAG101a/b chimeras accumulated to levels comparable with the native isoforms when expressed by Agroinfiltration, and also engaged into complexes with EDS1 (Figure 7d). Chimeras and native SAG101 isoforms were co-expressed with XopQ in *pss* mutant plants to test for functionality. SAG101a and the Nb-Ca chimeric protein did not show any activity. In contrast, the Na-Cb chimeric protein was able to restore XopQ-induced cell death, albeit to lesser extent than SAG101b (Figure 7e). These results suggest that main differences discriminating SAG101a/b and their immune competence reside in the C-terminal EP domain, and further supports that the EP domain surface is crucial for the function of EDS1 heterodimers in TNL signaling (Wagner et al., 2013; Bhandari et al., 2018). As the most probable mode of action of EDS1 complexes in immune signaling, we propose that further interaction partners might be recruited via the heterdodimeric EP domain surface. The newly established *N. benthamiana* system and mutant alleles will allow rapid verification of future hypotheses towards the elucidation of EDS1 immune functions.

**Figure 7:**
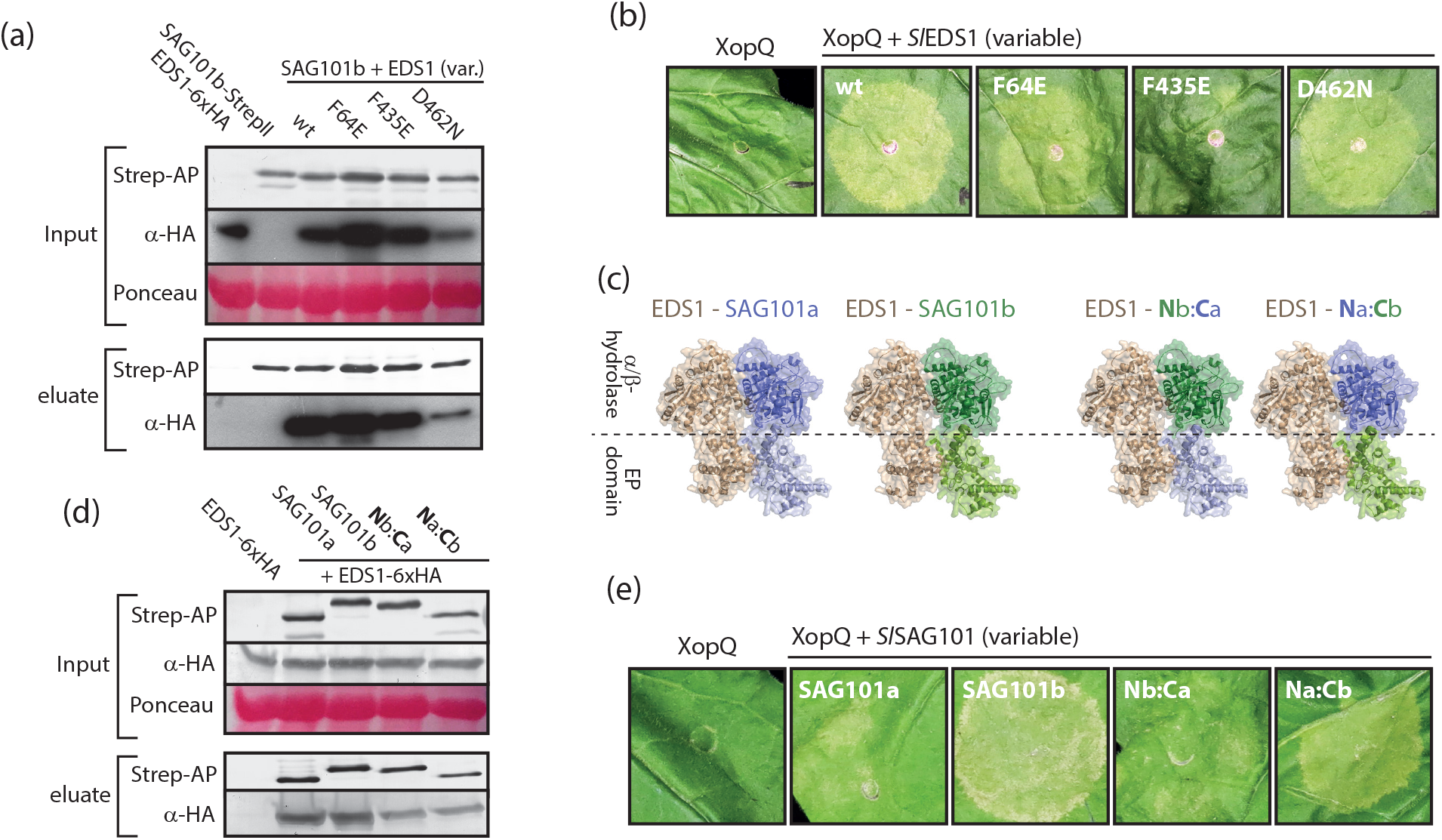
Identification of non-functional EDS1-SAG101b complex variants. a) Interaction of EDS1 variants with SAG101b. Indicated proteins were (co-) expressed in *N. benthamiana* by Agroinfiltration. Tissues were used 3 dpi for StrepII-purification. b) Immune activities of EDS1 variants. Indicated EDS1 variants (with C-terminal 6 x HA tag) were transiently co-expressed with XopQ-myc in *eds1* mutant plants by Agroinfiltration. Plant reactions were documented 7 dpi. c) Structural basis for S/SAG101a-S/SAG101b chimeric proteins. EDS1 and SAG101 both contain an N-terminal hydrolase-like and a C-terminal EP domain. In the heterodimer, an N-terminal interface is formed by the hydrolase-like domains, and a C-terminal interface by the EP domains. For chimeras, the N-terminus of SAG101b (aa 1-322) or SAG101a (aa 1-339) was fused with the C-terminus of SAG101a (aa 340-581) or SAG101b (aa 323-567), respectively. d) Heterocomplex formation by SAG101 chimeric proteins. SAG101 chimeras and native SAG101 isoforms (with a C-terminal StrepII tag) were co-expressed with EDS1-6xHA by Agroinfiltration. Tissues were used 3 dpi for StrepII-purification. e) Functionality of SAG101 chimeric proteins. Indicated proteins were expressed together with XopQ-myc in *pss* mutant plants by Agroinfiltration. Plant reactions were documented 7 dpi.

## Discussion

EDS1 is essential for signaling downstream of TNL-type immune receptors, and forms mutually exclusive heterodimeric complexes with PAD4 and SAG101 (Hu et al., 2005; Rietz et al., 2011; Wagner et al., 2013; Schultink et al., 2017). From analyses in the Arabidopsis system, immune functions were so far mainly accounted to EDS1-PAD4 (Feys et al., 2005; Wagner et al., 2013; Cui et al., 2017; Cui et al., 2018). Here, we show that an EDS1-SAG101 complex is necessary and sufficient for all tested TNL-dependent immune responses in *N. benthamiana,* while PAD4 does not appear to contribute to immunity (Figures 3, 4). This finding is particularly intriguing given that EDS1 and PAD4 are present in the genomes of all higher plants, while SAG101 is limited to those also encoding TNL-type immune receptors, strongly suggesting a functional link (Figure 1a; Wagner et al., 2013). It is thus tempting to speculate that EDS1-SAG101 might fulfil immune signaling functions in most, if not all, species containing TNL-type immune receptors outside the *Brassicaceae* family. Indeed, *Brassicaceae* PAD4 orthologues lack an insertion within the lipase-like domain, which is also absent in SAG101 orthologues (Wagner et al., 2013). Hence, PAD4 might have evolved by a unique path or mechanism in *Brassicaceae,* supporting the hypothesis that PAD4 immune signaling functions in this family might represent a notable exception. Future reverse genetic studies in additional species will clarify which EDS1-containing heterocomplexes function predominantly in TNL signaling.

Genome analysis revealed a duplication of SAG101 in most *Solanaceae* (Figure 1b). Absence of SAG101a in *Capsicum annuum* might indicate reduced selective forces towards its preservation. However, although both the *NbSAG101a2* and *NbSAG101b2* genes showed signs of pseudogenization in allotetraploid *N. benthamiana,* this was not observed for *SAG101a* in any of the remaining analyzed *Solanaceae* genomes. Also, different subcellular localizations were detected for SlSAG101 isoforms (Figures 2, S3), and SlSAG101a localized exclusively to the nucleus. This argues against SAG101a representing merely a duplicated gene, but rather supports distinct functions of individual isoforms. An additional EDS1-SAG101 complex might thus provide further fine-tuning of EDS1 activities in most *Solanaceae,* but we could not detect any contribution to TNL-mediated immune responses. We did not analyze functional relevance of EDS1 subcellular distribution in *N. benthamiana,* but nuclear localization was required and sufficient for several tested immune responses in Arabidopsis (Garcia et al., 2010; Stuttmann et al., 2016). It is worth noting that both *At*EDS1-*At*SAG101 and SlEDS1-SlSAG101a are restricted to nuclei and have only minor or no functions in immunity, while complexes required for TNL signaling (*At*EDS1-*At*PAD4, *S*/EDS1-*S*/SAG101b) are also distributed to the cytoplasm.

The *N. benthamiana* system established in this study features the key advantage of rapid and robust transient protein expression by Agroinfiltration. This was exploited for designing transient complementation assays for EDS1-SAG101 functional analyses based on induction of HR-like cell death by XopQ (Adlung et al., 2016; Schultink et al., 2017; Qi et al., 2018). We confirmed significance of results obtained in this highly simplified system by first disrupting EDS1-SAG101 complex formation (Figure 6). We mutagenized key residues within the interface, and show that higher order mutants containing multiple amino acid exchanges fail to function in immune signaling. Notably, several EDS1 and SAG101 variants, for which interaction was undetectable by co-purification (e.g. EDS1-TIVV, SAG101-FLL), could still function at least partially in cell death induction, indicating that low-level complex formation is sufficient for immune responses. Similarly, a previously described Arabidopsis PAD4-MLF variant (Wagner et al., 2013) deficient in complex formation fully complemented immune deficiency of a *pad4-1 sag101-3* double mutant line when tested (J. Stuttmann and J. Parker). Thus, extension of the interface analysis from EDS1 to its interaction partner SAG101 lends important support to the previous notion that complex formation is a prerequisite for immune signaling. But how do EDS1-PAD4 (in Arabidopsis) or EDS1-SAG101 (in *N. benthamiana*) contribute to immune signaling? Previous studies showed that a presumed EDS1 hydrolase activity is not required (Wagner et al., 2013).  Here, we identify several EDS1 alleles with reduced functionality, but retaining full protein stability and affinity to SAG101 (Figure 7). Most strikingly, the EDS1^F435D/E^ mutations fully abolish immune functions. Similarly, exchange in EDS1^F64E^ strongly affected EDS1 immune functions. These residues delimit upper and lower boundaries of an assumed cavity present on the surface of EDS1 heterodimers (Wagner et al., 2013; Bhandari et al., 2018). F64 might act as conserved gatekeeper, while perturbation of the C-terminal interface by F435E/D is expected to have more profound effects on overall topology of the EP domain assembly (Figure 7). A crucial role of the C-terminal EP domains for immune signaling is further supported by partial restoration of immune functions in SAG101a by grafting of the SAG101b C-terminus in SAG101^Na-Cb^ protein chimeras (Figure 7). We think these data are most compatible with recruitment of additional interaction partners via this surface. In agreement, residues bordering the assumed cavity and not involved in complex formation are also required for TNL signaling in Arabidopsis (Bhandari et al., 2018).

Interaction partner recruitment is further supported by results of our cross-species transfer of Roq1 and EDS1 family genes. NbRoq1 is functional in Arabidopsis (Figure 4b), and even immune receptors from evolutionary distant species maintain functionality when introduced to new plant lineages (Maekawa et al., 2012). In contrast, Arabidopsis EDS1-PAD4-SAG101 failed to function in *N. benthamiana* (Figure 5a). Similarly, tomato EDS1-PAD4 can fulfil immune functions in Arabidopsis but not *N. benthamiana,* while the opposite situation was observed for SlEDS1-SlSAG101b (Figures 5b, S6). We conclude that EDS1 complexes do not form a functional module in TNL signaling by themselves, but depend on additional factors, most likely interaction partners, co-evolving with the signaling-competent heterodimeric assembly in individual species. One expectation would be that mutant lines deficient in these factors phenocopy *eds1* lines. Although a number of proteins were previously reported to interact with EDS1 complexes (Arabidopsis Interactome Mapping, 2011; Bhattacharjee et al., 2011; Heidrich et al., 2011; Kim et al., 2012; Cui et al., 2018), mutant lines deficient in respective genes are not generally TNL signaling-deficient. Also, forward genetic screens in Arabidopsis failed to identify *eds1* phenocopies, suggesting genetic redundancy or a different molecular mode-of-action. However, a recent study in *N. benthamiana* identified the atypical CNL receptor NRG1 as another key component in TNL-mediated immunity (Qi et al., 2018). Similar to SAG101, CNLs of the NRG1 family are limited to those genomes also containing TNLs (Collier et al., 2011), and NRG1 may physically associate with EDS1 (Qi et al., 2018). Thus, proteins of the NRG1 class of helper CNLs may represent plausible candidate interaction partners, which could form a functional module with EDS1 complexes. Interestingly, *N. benthamiana nrg1* mutant plants still retain some competence to detect XopQ (via Roq1), and it was hypothesized that residual TNL signaling in these plants might be mediated by the ADR1 class of helper CNLs (Schultink et al., 2017; Qi et al., 2018). In Arabidopsis, there are three ADR1 homologs (ADR1-L1, −L2, −L3) and two NRG1 homologs (NRG1.1, NRG1.2) (Bonardi et al., 2011; Collier et al., 2011). Indeed, NRG1 homologs were recently reported to function redundantly in TNL-mediated immunity in Arabidopsis, and also ADR1s are required for several TNL-mediated immune responses (Dong et al., 2016; Castel et al., 2018; Wu et al., 2018). Helper CNLs of the NRG1 class are critical for function of most TNLs, but others require ADR1 class helpers or can signal via both pathways (Castel et al., 2018; Wu et al., 2018). While genetic redundancy explains failure to isolate respective mutant alleles by forward genetics, a physical association of NRG1 proteins with EDS1 as reported by Qi et al. (2018) will require further analysis. A similar interaction between *At*EDS1 and *At*NRG1.1 could be detected by Wu et al. (2018) only when using EDS1 as bait in co-IPs, and could results from stickiness of EDS1 in co-IP experiments (Wu et al., 2018). It should be noted that co-IPs in Qi et al. (2018) also show formation of NbEDS1 homodimers, which was included as positive control. We could not detect homodimerization of highly similar SlEDS1 in co-IPs, FRET-based interaction assays (Figures 2, S3) or by yeast two hybrid. The mechanistics underlying the functional relationships between TNLs, EDS1 complexes and helper CNLs thus remain a major question to pursue in future analyses.

We systematically compared *in planta* growth of avirulent (*Xcv* 85-10) and virulent (*Xcv ΔxopQ)* bacteria on wild type, *eds1, pss,* and *roq1 N. benthamiana* lines. In no case could we detect any basal resistance defect of *eds1* or *pss* lines (Figure 3). These findings are in agreement with previous reports from *N. benthamiana* and tomato, which failed to detect a basal resistance impairment in EDS1-deficient lines (Peart et al., 2002; Hu et al., 2005; Adlung et al., 2016; Schultink et al., 2017; Qi et al., 2018). We therefore think that EDS1 functions in basal resistance, supported by enhanced growth of several bacterial, fungal or oomycete isolates (e.g. Falk et al., 1999; Lipka et al., 2005; Rietz et al., 2011; Schon et al., 2013) on *eds1* deficient Arabidopsis lines, most likely result from loss of TNL-mediated ETI rather than an independent function of EDS1. This is in line with expression of basal resistance by the term “MTI + weak ETI – effector-triggered susceptibility” (Jones and Dangl, 2006), and the TNL-mediated component of “weak ETI” being abolished in *eds1* lines. We therefore favor the model that EDS1 complexes fulfil one single function tightly linked to TNL signaling in plant immunity, and that any differential effects of particular alleles on basal vs. TNL-mediated immunity or cell death can be ascribed to detection thresholds of experimental systems. Accordingly, the functions of EDS1-PAD4 in organisms lacking TNLs remain largely unexplored, but go along with perfect conservation of the hydrolase catalytic triad, dispensable for TNL signaling, in both proteins (Pegadaraju et al., 2005; Pegadaraju et al., 2007; Louis et al., 2012; Wagner et al., 2013; Chen et al., 2018). In contrast, EDS1-SAG101 (or, in exceptions, EDS1-PAD4) were recruited to TNL-signaling in the dicot plant lineage. TNL signaling depends on a non-catalytic mechanism, and an intact catalytic triad was so far not detected in SAG101 homologs. This sector of the immune network was lost e.g. in the order Lamiales and columbine *(Aquilegia coerulea),* correlating with the concurrent loss of SAG101 and NRG1 (Collier et al., 2011; Jacob et al., 2013; Wagner et al., 2013). From our data, we propose that immune signaling functions of EDS1-SAG101 in *N. benthamiana* rely on the recruitment of further interaction partners via the EP domain. The *N. benthamiana* system established in this study will facilitate future functional analyses towards a molecular understanding of TNL signaling and EDS1 functions.

## Material and Methods

### Plant material, growth conditions, bacterial strains and infection assays

*N. benthamiana* wild type plants and the published *eds1a-1* and *pad4-1* single and *eds1a-1 pad4-1* double mutant lines were used (Ordon et al., 2017). *N. benthamiana* plants were cultivated in a greenhouse with 16 h light period, 60 % relative humidity at 24/20 °C (day/night). *A. thaliana* wild type accession Columbia and the previously published *eds1-2 pad4-1* double and *eds1-2 pad4-1 sag101-1* triple mutant lines were used (Feys et al., 2005; Wagner et al., 2013). Arabidopsis plants were grown under short day conditions at 23/21 °C and with 60 % relative humidity or in a greenhouse under long day conditions for seed set. For bacterial growth assays, the *Xcv* strain 85-10 (Thieme et al., 2005) and the *ΔxopQ* mutant (Adlung et al., 2016) were syringe-infiltrated at an OD_600_ = 0.0004, leaf discs harvested with a cork borer at different time points, disrupted in 10 mM MgCl_2_ using a bead mill, and bacterial titers determined by plating dilution series. For each time point and strain, samples were taken from at least four independent leaves, and treated as biological replicates. Bacterial growth assays were repeated at least three times with similar results. For type III system-dependent protein translocation via *Pseudomonas fluorescence,* a previously described derivative of the “EtHAn” strain (Thomas et al., 2009) containing a plasmid for translocation of XopQ fused to a secretion signal of AvrRpt2 was used (Gantner et al., 2018). *Hyaloperonospora arabidopsidis* isolate Cala2 was used for infection of Arabidopsis plants, and infections were done as previously described (Wagner et al., 2013). True leaves were stained with Trypan Blue 7 dpi, and representative micrographs are shown.

### Phylogenetic analyses

Genomes as indicated in Supplemental Table 1 were mined for *EDS1* family genes by tBLASTn using tomato proteins as query. Gene models were examined or assigned using fgenesh+ (Solovyev, 2007) and multiple sequence alignments. Phylogenetic tree construction was done using the phylogeny.fr platform (Dereeper et al., 2008) with standard settings.

### *Agrobacterium*-mediated expression, StrepII-purification and immunodetection

For transient *Agrobacterium*-mediated expression of proteins in *N. benthamiana* (“Agroinfiltration”), plate-grown bacteria were resuspended in Agrobacterium Infiltration Medium (AIM; 10 mM MES pH 5.8, 10 mM MgCl_2_). Single strains were infiltrated at an OD_600_ = 0.6. For co-expression, an OD_600_ = 0.4 for each strain was used. All constructs for expression of proteins in *N. benthamiana* contained the 35S promoter. EDS1, PAD4, SAG101 or variants were co-expressed with XopQ-myc in reconstitution assays. For immunodetection of proteins in support of reconstitution assays and for co-purification, proteins were expressed without XopQ to avoid interference due to the negative effect of XopQ recognition on *Agrobacterium*-mediated protein expression (Adlung and Bonas, 2017), but using the same *N. benthamiana* genetic background as in respective reconstitution experiments. Leaf tissue was ground in liquid nitrogen, powder resuspended in Laemmli buffer and proteins denatured by boiling prior to SDS-PAGE for immunodetection. For StrepII-purifications, 1 g of leaf tissue was ground in liquid nitrogen and the leaf powder resuspended in 2.5 ml of extraction buffer (50 mM Tris pH8, 150 mM NaCl, 5 mm EDTA, 5mM EGTA, 10 mM DTT, 0,1 % Triton X-100). Suspensions were cleared by centrifugation and supernatants passed through a 0.45 μm syringe filter. 2 ml of cleared extracts were incubated with 120 μl of Strep-Tactin high-capacity matrix (IBA) for 20 min at 4°C on a rotary wheel. The matrix was washed several times with extraction buffer prior to elution of proteins by boiling with 100 μl of Laemmli buffer. Proteins were resolved by SDS-PAGE and transferred to a nitrocellulose membrane (GE Healthcare). StrepII-tagged proteins were detected using Strep-Tactin AP conjugate (IBA) or a mouse monoclonal StrepII antibody (Sigma). Further primary antibodies used were α-mCherry (Abcam, ab167453,), mouse monoclonal α-GFP and α-c-myc, rat α-HA (all from Roche), and α-FLAG (Sigma). Secondary antibodies were coupled to horseradish peroxidase (HRP, GE Healthcare) or alkaline phosphatase (AP, Sigma).

### Plant transformation and genome editing

Arabidopsis plants were transformed as previously described (Logemann et al., 2006). For transformation of *N. benthamiana,* leaves of greenhouse-grown plants were surface sterilized, cut and co-cultivated with Agrobacteria containing Cas9/sgRNA constructs. Explants were surface-sterilized, and transgenic plants regenerated. A detailed protocol is provided as an online resource (dx.doi.org/10.17504/protocols.io.sbaeaie). Details on constructs, sgRNAs for editing of *N. benthamiana SAG101* isoforms and *Roq1* and generated mutant alleles are provided in Figure S4. Primary transformants (T0 plants) were tested phenotypically by challenge inoculation with XopQ-translocating *Pseudomonas fluorescence* bacteria, and screened by PCR. A transgene-free *pad4-1 sag101a-1 sag101b-1 (pss)* triple mutant was isolated from a segregating T_1_ population by PCR screening, and crossed to wild type for isolation of the *pad4-1 sag101b-1* double mutant line. Homozygous, non-transgenic seed lots were used for experiments. In case of the *roq1* mutant line, a T_1_ population segregating for two different disruptive alleles (*roq1-3* and *roq1-4;* Figure S4), was used for infection assays.

### Life Cell Imaging and FRET-APB Analysis

Images were taken on a Zeiss LSM780 laser scanning microscope. For imaging of GFP and mCherry, fluorophores were excited with 488 nm and 561 nm laser lines, and emission detected at 493-556 nm and 597-636 nm, respectively. For intensity-based FRET (FRET-APB), mCherry was bleached using the 561 nm laser at 100% intensity, and GFP fluorescence measured pre- and post-bleach. At least 15 measurements per donor/acceptor combination were done per experiment, and data was reproduced in four independent repetitions.

### Molecular cloning and yeast two hybrid interaction assays

Constructs were generated by Golden Gate (Engler et al., 2008) and Gateway (Thermo Fisher Scientific; as according to manufacturer’s instructions) cloning. Golden Gate reactions with either *BsaI* or *BpiI* were performed using 20-40 fmol of each DNA module and cycling between 37 °C and 16 °C, as previously described (Weber et al., 2011). DNA modules of the MoClo Plant Toolkit, Plant Parts I (Engler et al., 2014) and Plant Parts II (Gantner et al., 2018) collections were used. Novel Level 0 modules were generated as previously described, and restriction sites eliminated by site-directed mutagenesis or overlapping PCR products (Engler et al., 2008; Engler et al., 2014). Details on generated constructs and oligonucleotides used for cloning are provided in Table S3. Previously described (Gantner et al., 2018) Golden Gate-compatible or Gateway-converted derivatives of pGADT7 and pGBKT7 (Clontech) were used for yeast two hybrid assays. Respective constructs were transformed in yeast strain PJ69-4a by standard procedures (Gietz and Schiestl, 2007). Plate-selected co-transformants were cultivated in liquid SD media for 48h, dilution series prepared and plated on selective media using a multipipette. Extraction of proteins for immunodetection was performed as previously described (Kushnirov, 2000).

### Gene expression analysis

Tomato RNA sequencing data was accessed using the TomExpress portal (Zouine et al., 2017; http://tomexpress.toulouse.inra.fr/). Data was visualized as a normalized expression heatmap using Spearman representation. Expression values for different conditions were added manually. For gene expression analyses in *N. benthamiana,* plants were syringe infiltrated with *Xcv* bacteria at an OD_600_ = 0.02 in 10 mM MgCl_2_, or mock-infiltrated. RNA was extracted by a standard protocol using TRIzol™ reagent (ambion; Fisher Scientific). Briefly, two leaf discs (diameter 9 mm) were frozen in liquid nitrogen and tissues disrupted using Zirkonia beads (Carl Roth N039.1) and a bead mill. RNA was extracted with 1 ml TRIzol, 100 μl of bromochloropropane were added for phase separation, and RNA was precipitated, washed, dried and resuspended in 40 μl H_2_O. The Reverse Transcriptase Core Kit was used for cDNA synthesis, and the Takyon No ROX SYBR 2X MasterMix blue dTTP qPCR Kit (both Eurogentech) for quantitative real time PCR using a CFX96 detection system (Bio-Rad). The previously described reference genes *Protein Phosphatase 2A (PP2A)* and *Elongation Factor 1-α (EF1α)* were used for data normalization (Liu et al., 2012) with similar results, and data from normalization to *PP2A* is shown. Primers used for quantitative real PCR are listed in Table S4. All primers had efficiencies of 90-105%, as evaluated by dilution series.

### Protein Modeling

Structural models of *Solanum lycopersicum* EDS1-SAG101a and EDS1-SAG101b complexes were modeled using the structure of the *Arabidopsis thaliana* EDS1-SAG101 as template (PDB: 4NFU) (PDB: 4NFU; Wagner et al., 2013). Sequences of SlSAG101a and SlSAG101b share 38% and 36% sequence identity with AtSAG101, respectively, while *S*/EDS1 and *At*EDS1 share 40%. All three sequence-template pairs could therefore be confidently aligned using hhpred algorithm (Zimmermann et al., 2018) and structural models were generated and relaxed based on these alignments using rosettaCM (Song et al., 2013) with limited need for manual re-alignment in the regions with insertions. Sequence conservation was calculated using the rate4site algorithm (Pupko et al., 2002) and mapped at the surface of structural models using pymol (The PyMOL Molecular Graphics System, Version 2.0 Schrödinger, LLC). Structural models and analysis thereof are provided in Appendix S1.

## Acknowledgements

We acknowledge Bianca Rosinsky for taking care of plant growth facilities and growing plants. Ulla Bonas is acknowledged for generous support. Martin Schattat is acknowledged for assistance with FRET experiments. This work was funded by GRC grant STU 642-1/1 (Deutsche Forschungsgemeinschaft, DFG) and seed funding by the CRC 648 (DFG) to Johannes Stuttmann.

---

**Supplemental Figure S1:**
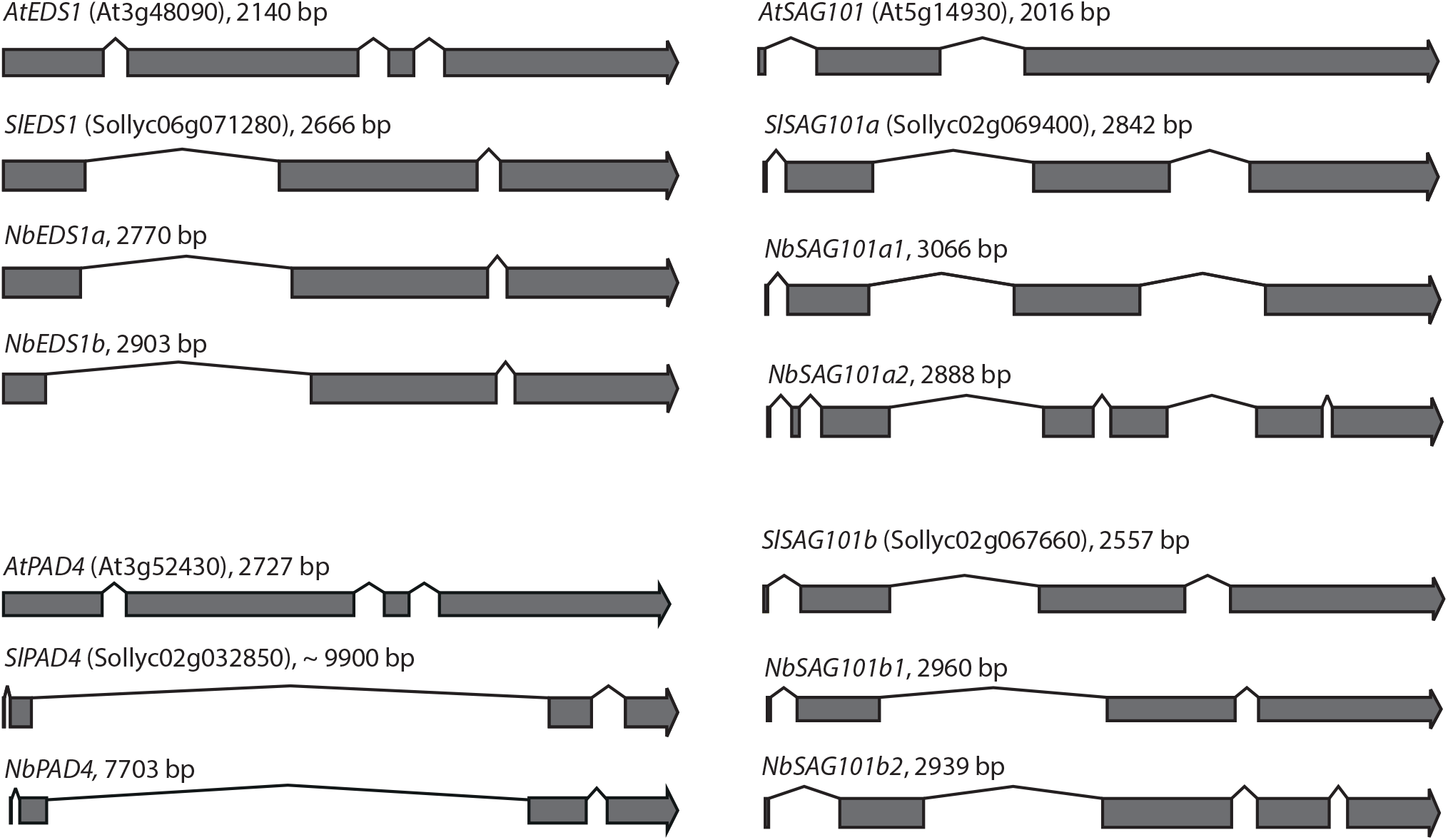
*EDS1* family gene models from Arabidopsis, tomato and *Nicotiana benthamiana* Putative *EDS1, PAD4* and *SAG101* homologs were detected as described in materials and methods. Gene models (*Nicotiana benthamiana*) were predicted using fgenesh+ and the corresponding tomato proteins as support. Sequence details including annotations are provided in Table S1.

**Supplemental Figure S2:**
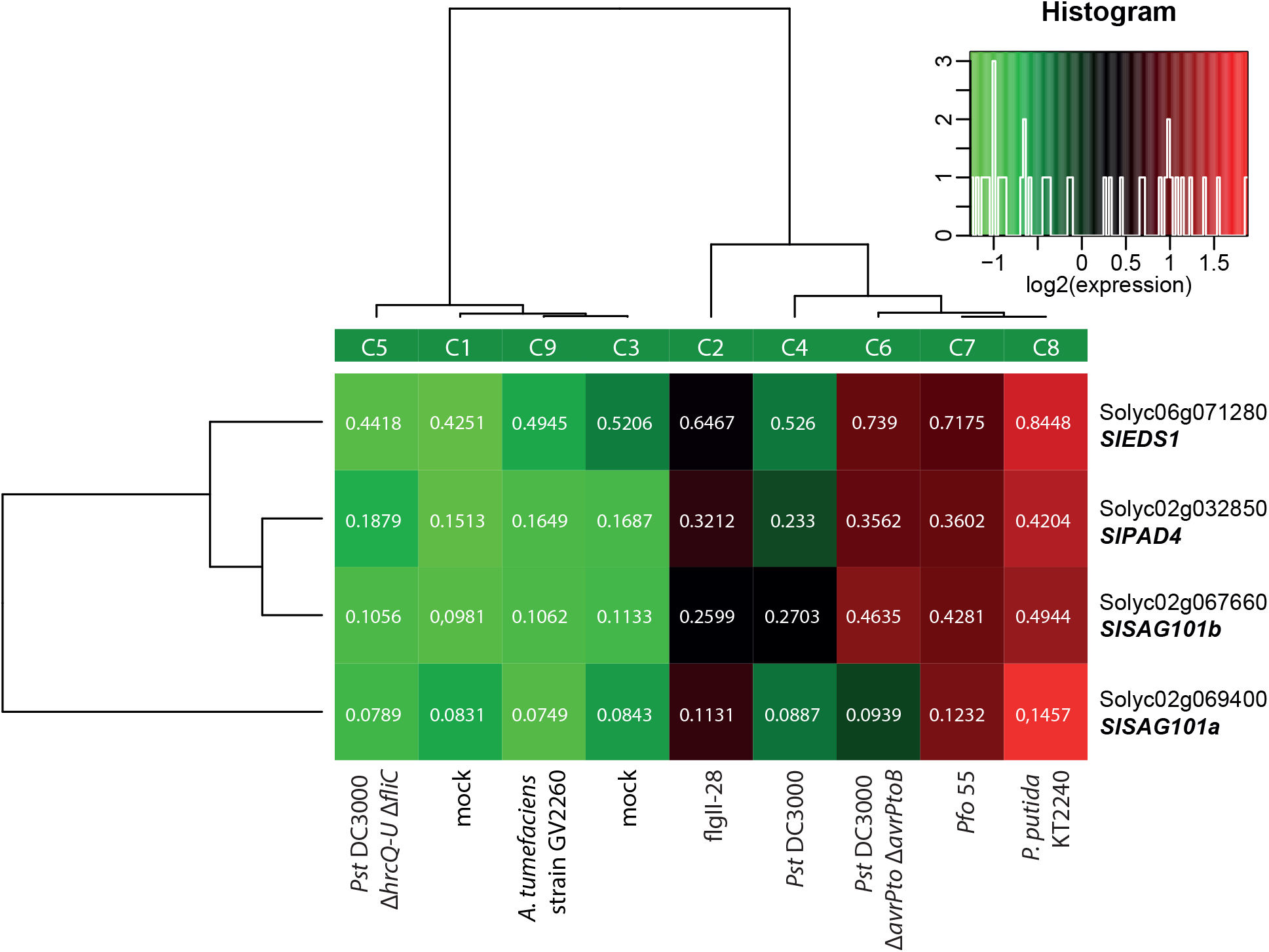
Expression of tomato *EDS1* family genes. A public RNAseq dataset from treatment of tomato “Rio Grande” plants with different MTI-inducers (Rosli et al., 2013) was analyzed for expression of *EDS1* family genes using the TomExpress portal (Zouine et al., 2017). Hierarchical clustering using Spearman distance (output from TomExpress) is shown.

**Supplemental Figure S3:**
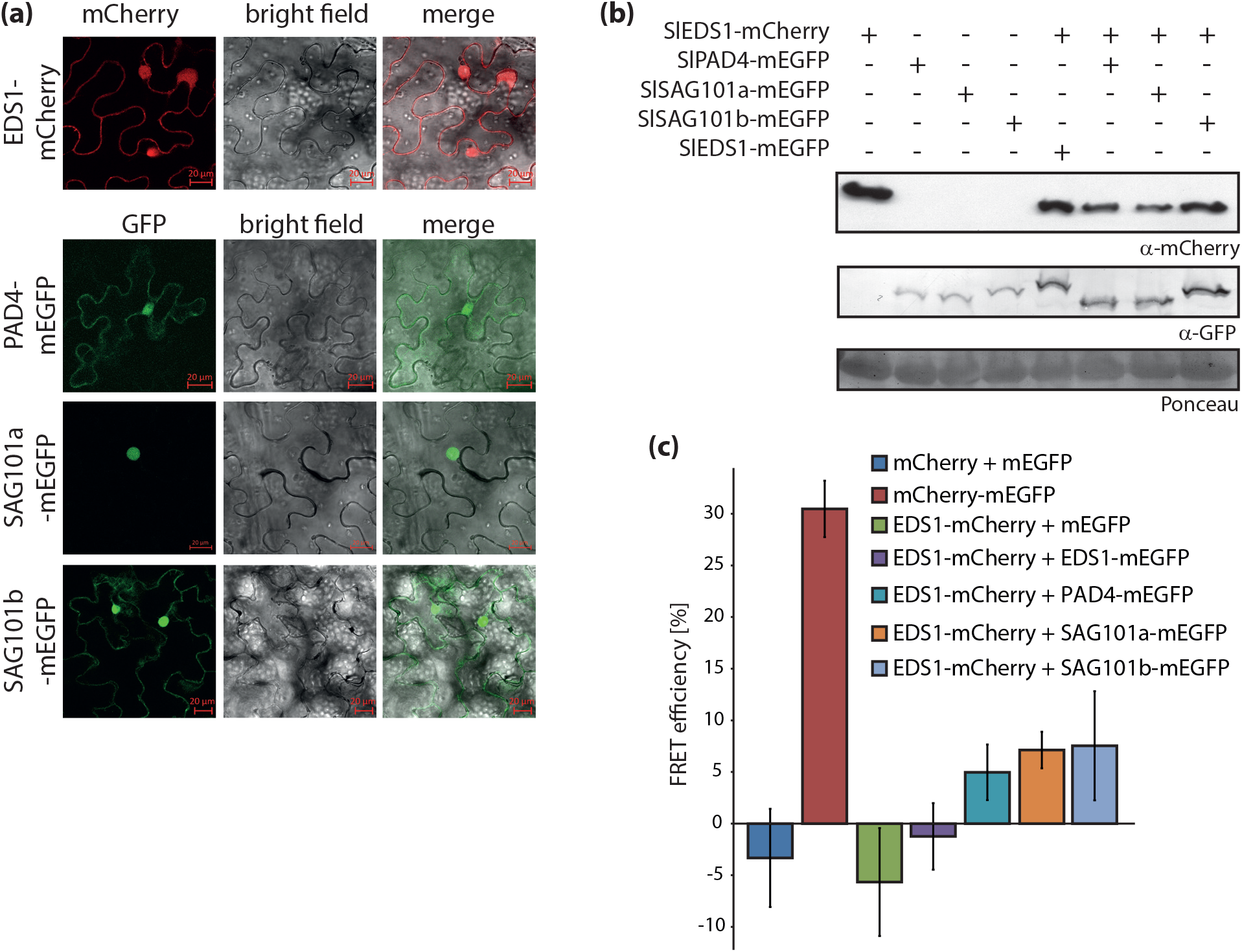
Localization and complex formation of tomato EDS1 proteins. a) Localization of tomato EDS1, PAD4 and SAG101a/b isoforms when expressed singly. Extended data supporting Figure 2a. b) Integrity of EDS1, PAD4 and SAG101 fluorophore fusions when expressed singly or in combination. Extended data supporting Figures 2a, S3a and S3c. c) Formation of complexes between tomato EDS1 family proteins in living cells as measured by FRET-APB (intensity-based FRET; n ≥ 30, standard deviation).

**Supplemental Figure S4:**
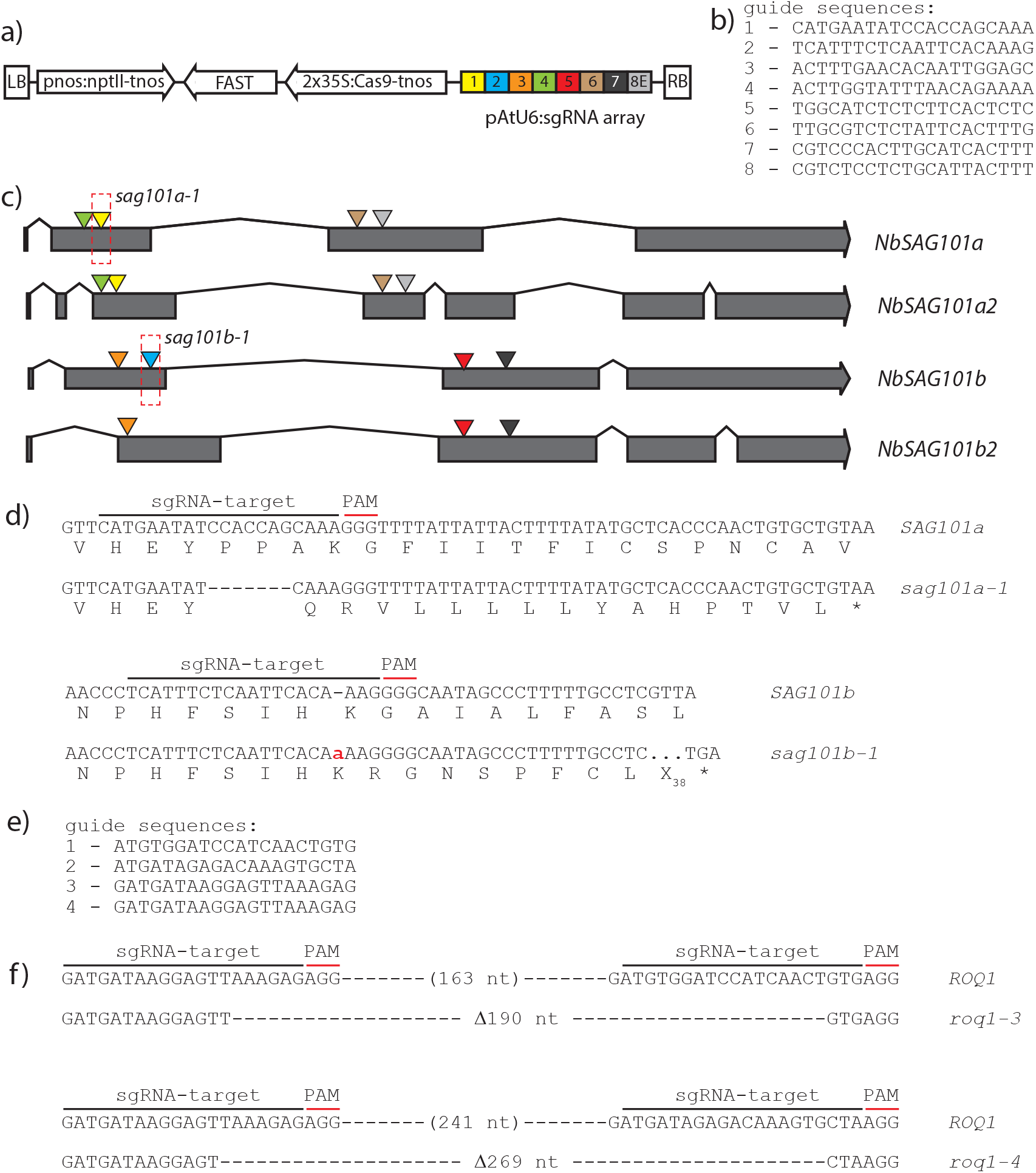
Generation of mutant lines by genome editing. a) Scheme of the T-DNA construct used for editing *SAG101* genes in *N. benthamiana.* Construct is based on pDGE160 (Ordon et al., 2017). b) Guide sequences incorporated in the sgRNA array of the construct shown in a). c) Position of target sites within *SAG101* gene models. The color code corresponds to panel a). d) Details on *sag101a-1* and *sag101b-1* alleles generated by genome editing. e) Guide sequences used for editing of the *Roq1* gene. A construct similar to that in a), but based on a different pDGE recipient vector (pDGE311), was used. f) Molecular details on *roq1-3* and *roq1-4* alleles generated by genome editing.

**Supplemental Figure S5:**
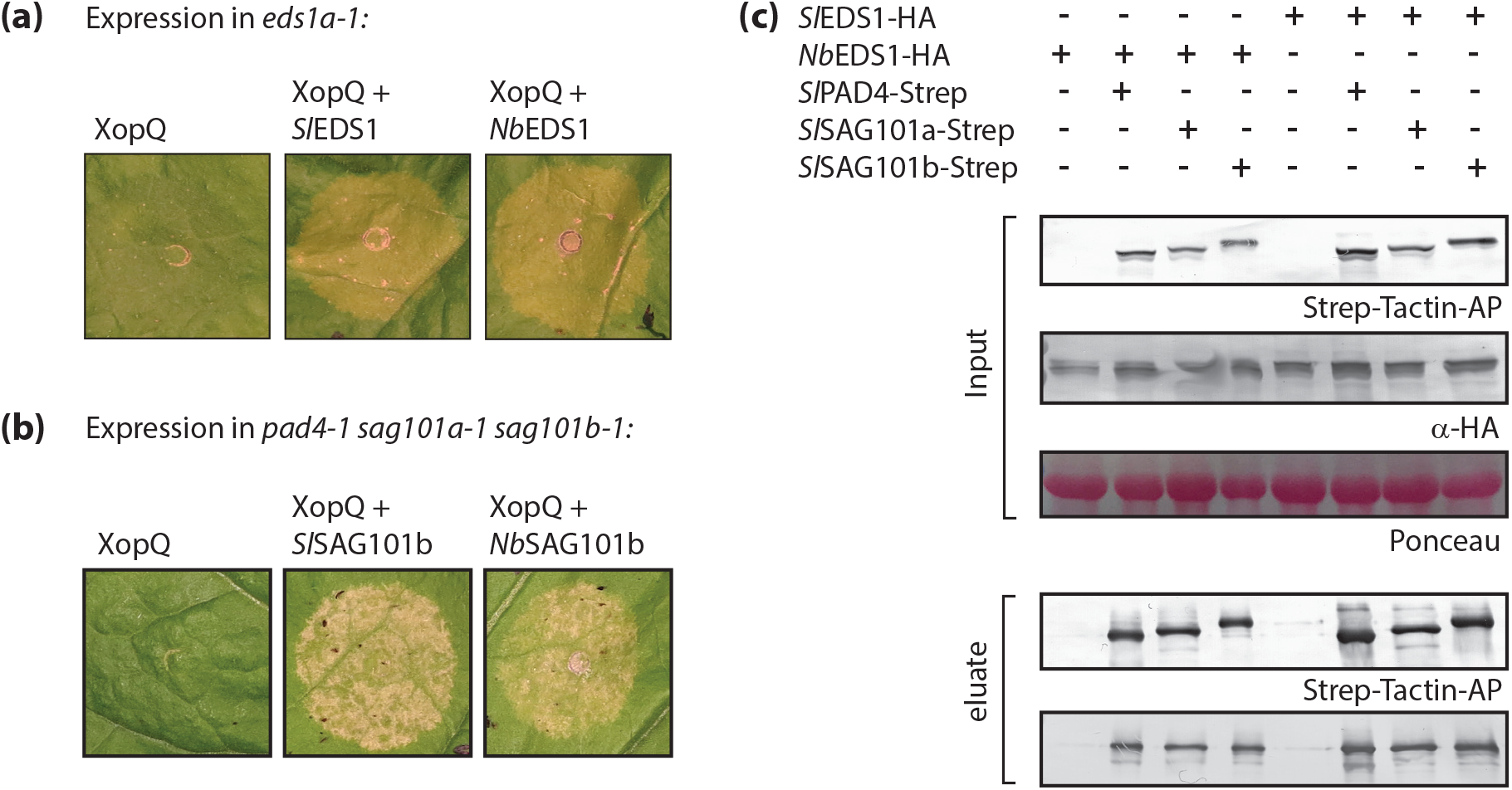
Functional comparison of EDS1 and SAG101b from *N. benthamiana* and *S. lycopersicum* for XopQ-induced cell death. a) Restoration of XopQ-induced cell death by co-expression of *S*/EDS1 and *Nb*EDS1. Indicated proteins were (co-) expressed in *eds1* mutant plants, and HR development was documented 6 dpi. b) As in (a), but *S*/SAG101b and *Nb*SAG101b were co-expressed together with XopQ in *pad4 sag101a sag101b* triple mutant plants. c) Complex formation between *Nb*EDS1 and *S*/PAD4, *S*/SAG101a and *S*/SAG101b. Indicated proteins were (co-) expressed in *N. benthamiana* by Agroinfiltration. Tissues were used 3 dpi for StrepII-purifìcation.

**Supplemental Figure S6:**
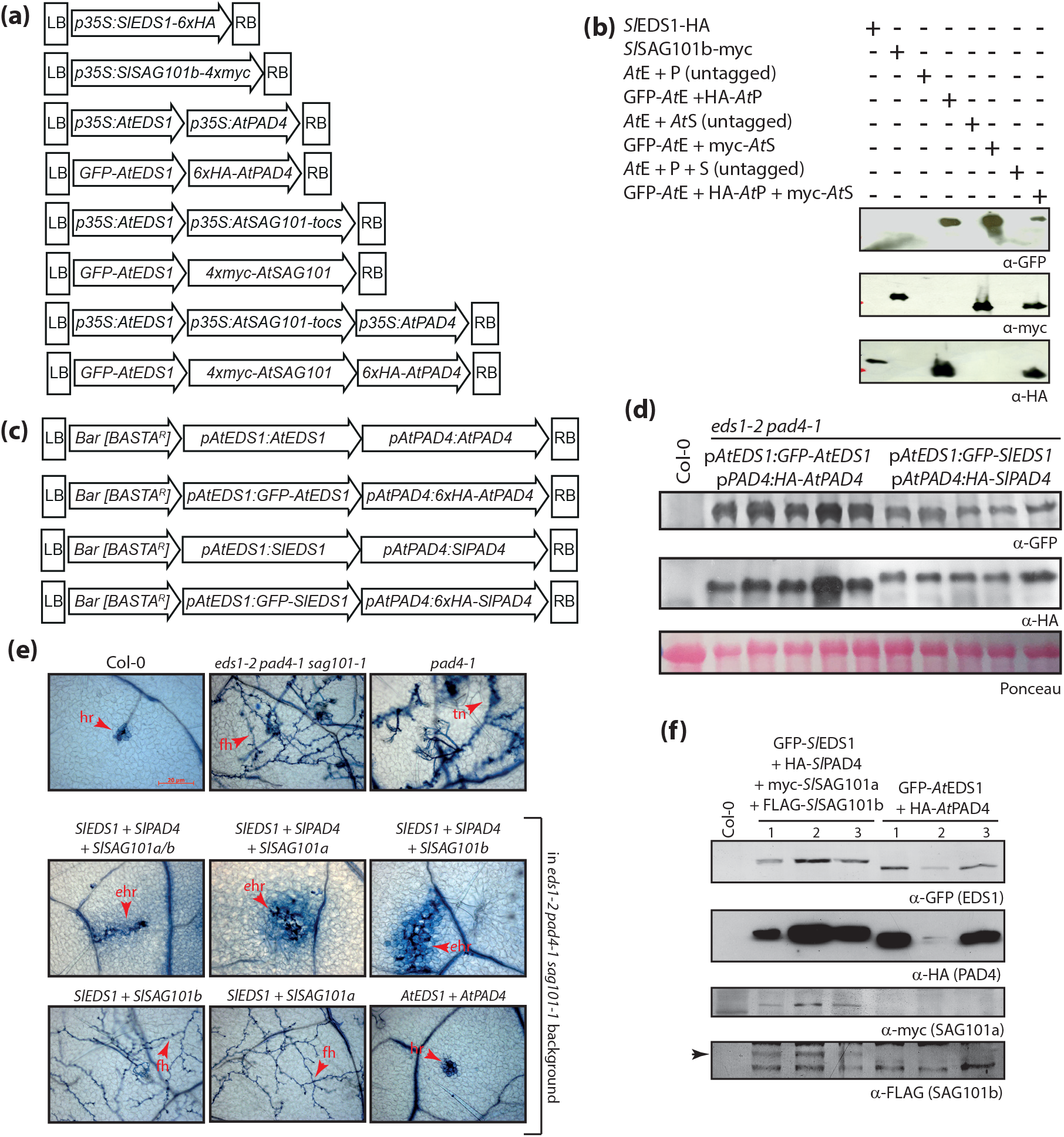
Cross-species transfer of *EDS1*-family genes. a) Schematic representation of T-DNA constructs used for *Agrobacterium*-mediated transient expression of *EDS1* family genes (from Arabidopsis or tomato) in *N. benthamiana.* Extended data to Figure 5a. b) Expression of Arabidopsis proteins in *N. benthamiana.* Extended data to Figure 5a. E – EDS1, P – PAD4, S – SAG101. Total extracts were prepared from infiltrated leaf sections 3 dpi for protein detection. c) Schematic representation of T-DNA constructs used for Arabidopsis transformation. d) Immunodetection of transgenic protein expression. Three week-old BASTA-resistant T_2_ plants of individual families were pooled for protein extraction. Ponceau staining is shown as loading control. e) Functionality of tomato EDS1 family proteins in Arabidopsis. Indicated combinations of tomato genes under control of the corresponding Arabidopsis promoter elements were expressed (with or without an epitope tag) in the *eds1-2 pad4-1 sag101-1* triple mutant background. Constructs were of similar architecture as before (Figure S6c), but contained the FAST marker. T lines from transformation of constructs without epitope tags were used for infection assays: Transformed T seeds were selected by FAST seed fluorescence, three week-old plants used for infection with *Hpa* isolate Cala2, and tissues stained with Trypan Blue 7 dpi. Similarly selected plants expressing tagged proteins were used for immunodetection (Figure S6f). hr – hypersensitive response; fh – free hyphae; tn – trailing necrosis; ehr – expanded hypersensitive response. f) Immunodetection of tomato EDS1 family proteins in transgenic Arabidopsis. Each lane represents an individual T_4_ plant.

**Supplemental Figure S7:**
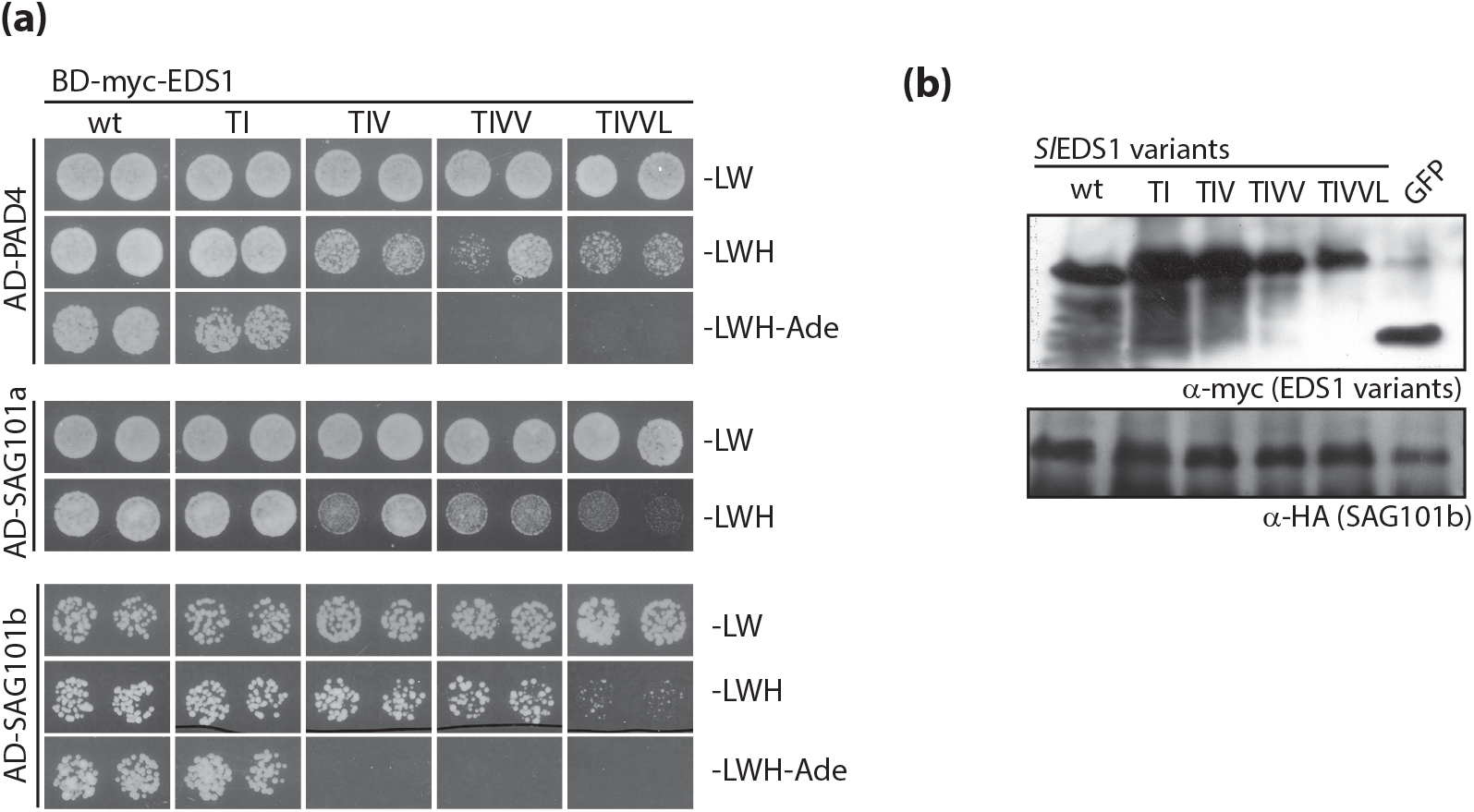
Heterocomplex formation by EDS1 variants in a yeast two hybrid system. a) Yeast two hybrid interaction assay. pGADT7 and pGBKT7 derivatives coding for the indicated protein fusions were co-transformed into yeast strain PJ69-4a. Two independent transformants were grown in dilution series on media lacking leucine and tryptophan (growth), or additionally lacking histidine (interaction; low stringency reporter) or histidine and adenine (interaction; high stringency reporter. Yeast plates were incubated at 30°C for 3d prior to documentation. A higher dilution is shown for the SAG101b-EDS1 interaction assay. b) Immunodetection of fusion proteins expressed in yeast transformants from a).

**Supplemental Figure S8:**
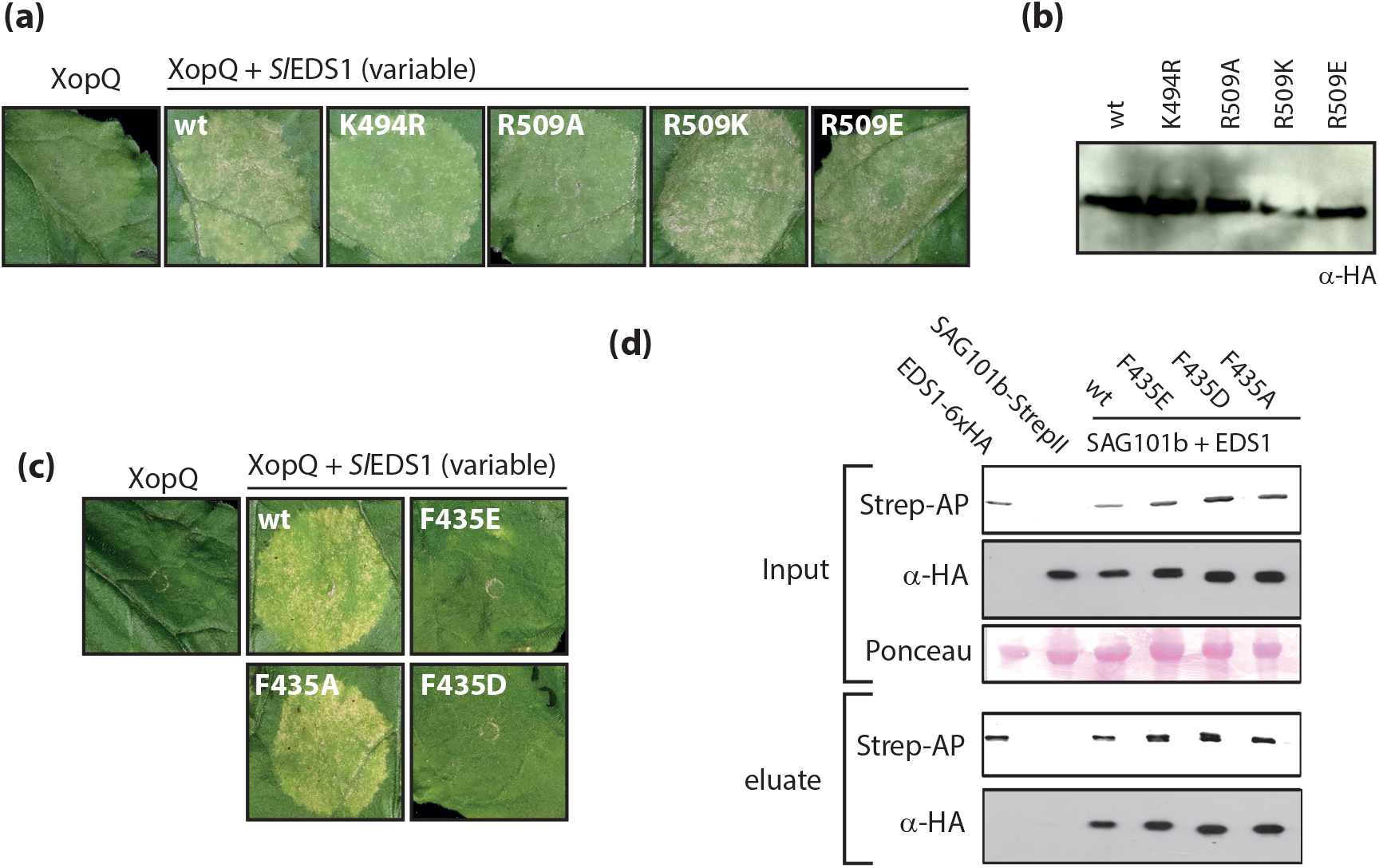
Immune competence of further EDS1 variants. a) Functionality of EDS1 variants carrying exchanges in positively charged residues lining a cavity on the EDS1 surface. Indicated variants were co-expressed with XopQ-myc in *eds1* mutant plants, and plant reactions documented 7 dpi. b) Immunodetection of EDS1 variants used in a). c) Functionality of EDS1 variants carrying exchanges in C-terminal heterocomplex interface. Indicated variants were co-expressed with XopQ-myc in *eds1* mutant plants, and plant reactions documented 7 dpi. d) Immunodetection of EDS1 variants used in c).

## References

Aarts, N., Metz, M., Holub, E., Staskawicz, B.J., Daniels, M.J., and Parker, J.E. (1998). Different requirements for *EDS1* and *NDR1* by disease resistance genes define at least two *R* gene-mediated signaling pathways in *Arabidopsis*. Proceedings of the National Academy of Sciences of the United States of America 95, 10306–10311.

Adlung, N., and Bonas, U. (2017). Dissecting virulence function from recognition: cell death suppression in Nicotiana benthamiana by XopQ/HopQ1-family effectors relies on EDS1-dependent immunity. Plant J 91, 430–442.

Adlung, N., Prochaska, H., Thieme, S., Banik, A., Bluher, D., John, P., Nagel, O., Schulze, S., Gantner, J., Delker, C., Stuttmann, J., and Bonas, U. (2016). Non-host Resistance Induced by the Xanthomonas Effector XopQ Is Widespread within the Genus Nicotiana and Functionally Depends on EDS1. Front Plant Sci 7, 1796.

Arabidopsis Interactome Mapping, C. (2011). Evidence for network evolution in an Arabidopsis interactome map. Science (New York, N.Y 333, 601–607.

Baggs, E., Dagdas, G., and Krasileva, K.V. (2017). NLR diversity, helpers and integrated domains: making sense of the NLR IDentity. Current opinion in plant biology 38, 59–67.

Bent, A.F., Kunkel, B.N., Dahlbeck, D., Brown, K.L., Schmidt, R., Giraudat, J., Leung, J., and Staskawicz, B.J. (1994). *RPS2* of *Arabidopsis thaliana*: A leucine-rich repeat class of plant disease resistance genes. Science (New York, N.Y 265, 1856–1860.

Bentham, A., Burdett, H., Anderson, P.A., Williams, S.J., and Kobe, B. (2017). Animal NLRs provide structural insights into plant NLR function. Ann Bot 119, 827–702.

Bernoux, M., Ve, T., Williams, S., Warren, C., Hatters, D., Valkov, E., Zhang, X., Ellis, J.G., Kobe, B., and Dodds, P.N. (2011). Structural and functional analysis of a plant resistance protein TIR domain reveals interfaces for self-association, signaling, and autoregulation. Cell Host Microbe 9, 200–211.

Bernoux, M., Burdett, H., Williams, S.J., Zhang, X., Chen, C., Newell, K., Lawrence, G.J., Kobe, B., Ellis, J.G., Anderson, P.A., and Dodds, P.N. (2016). Comparative Analysis of the Flax Immune Receptors L6 and L7 Suggests an Equilibrium-Based Switch Activation Model. The Plant cell 28, 146–159.

Bhandari, D.D., Lapin, D., Kracher, B., vonBorn, P., Bautor, J., Niefind, K., and Parker, J.E. (2018). An EDS1 EP-domain surface mediating timely transcriptional reprogramming of immunity genes. bioRxiv.

Bhattacharjee, S., Halane, M.K., Kim, S.H., and Gassmann, W. (2011). Pathogen effectors target Arabidopsis EDS1 and alter its interactions with immune regulators. Science (New York, N.Y 334, 1405–1408.

Bonardi, V., Tang, S., Stallmann, A., Roberts, M., Cherkis, K., and Dangl, J.L. (2011). Expanded functions for a family of plant intracellular immune receptors beyond specific recognition of pathogen effectors. Proceedings of the National Academy of Sciences of the United States of America 108, 16463–16468.

Burch-Smith, T.M., Schiff, M., Caplan, J.L., Tsao, J., Czymmek, K., and Dinesh-Kumar, S.P. (2007). A novel role for the TIR domain in association with pathogen-derived elicitors. PLoS biology 5, e68.

Büttner, D. (2016). Behind the lines-actions of bacterial type III effector proteins in plant cells. FEMS Microbiol Rev.

Castel, B., Ngou, P.M., Cevik, V., Redkar, A., Kim, D.S., Yang, Y., Ding, P., and Jones, J.D.G. (2018). Diverse NLR immune receptors activate defence via the RPW8-NLR NRG1. New Phytol.

Century, K.S., Shapiro, A.D., Repetti, P.P., Dahlbeck, D., Holub, E., and Staskawicz, B.J. (1997). *NDR1*, a pathogen-induced component required for *Arabidopsis* disease resistance. Science (New York, N.Y 278, 1963–1965.

Cesari, S. (2018). Multiple strategies for pathogen perception by plant immune receptors. New Phytol 219, 17–24.

Chakravarthy, S., Velasquez, A.C., Ekengren, S.K., Collmer, A., and Martin, G.B. (2010). Identification of Nicotiana benthamiana genes involved in pathogen-associated molecular pattern-triggered immunity. Mol Plant Microbe Interact 23, 715–726.

Chen, G., Wei, B., Li, G., Gong, C., Fan, R., and Zhang, X. (2018). TaEDS1 genes positively regulate resistance to powdery mildew in wheat. Plant molecular biology 96, 607–625.

Collier, S.M., Hamel, L.P., and Moffett, P. (2011). Cell death mediated by the N-terminal domains of a unique and highly conserved class of NB-LRR protein. Mol Plant Microbe Interact 24, 918–931.

Cui, H., Tsuda, K., and Parker, J.E. (2015). Effector-triggered immunity: from pathogen perception to robust defense. Annu Rev Plant Biol 66, 487–511.

Cui, H., Gobbato, E., Kracher, B., Qiu, J., Bautor, J., and Parker, J.E. (2017). A core function of EDS1 with PAD4 is to protect the salicylic acid defense sector in Arabidopsis immunity. New Phytol 213, 1802–1817.

Cui, H., Qiu, J., Zhou, Y., Bhandari, D.D., Zhao, C., Bautor, J., and Parker, J.E. (2018). Antagonism of Transcription Factor MYC2 by EDS1/PAD4 Complexes Bolsters Salicylic Acid Defense in Arabidopsis Effector-Triggered Immunity. Mol Plant 11, 1053–1066.

Dereeper, A., Guignon, V., Blanc, G., Audic, S., Buffet, S., Chevenet, F., Dufayard, J.-F., Guindon, S., Lefort, V., Lescot, M., Claverie, J.-M., and Gascuel, O. (2008). Phylogeny.fr: robust phylogenetic analysis for the non-specialist. Nucleic Acids Research 36, 465–469.

Dong, O.X., Tong, M., Bonardi, V., El Kasmi, F., Woloshen, V., Wunsch, L.K., Dangl, J.L., and Li, X. (2016). TNL-mediated immunity in Arabidopsis requires complex regulation of the redundant ADR1 gene family. New Phytol 210, 960–973.

Engler, C., Kandzia, R., and Marillonnet, S. (2008). A one pot, one step, precision cloning method with high throughput capability. PLoS ONE 3, e3647.

Engler, C., Youles, M., Gruetzner, R., Ehnert, T.M., Werner, S., Jones, J.D., Patron, N.J., and Marillonnet, S. (2014). A Golden Gate Modular Cloning Toolbox for Plants. ACS synthetic biology.

Falk, A., Feys, B.J., Frost, L.N., Jones, J.D., Daniels, M.J., and Parker, J.E. (1999). EDS1, an essential component of *R* gene-mediated disease resistance in Arabidopsis has homology to eukaryotic lipases. Proceedings of the National Academy of Sciences of the United States of America 96, 3292–3297.

Feys, B.J., Moisan, L.J., Newman, M.A., and Parker, J.E. (2001). Direct interaction between the Arabidopsis disease resistance signaling proteins, EDS1 and PAD4. EMBO Journal 20, 5400–5411.

Feys, B.J., Wiermer, M., Bhat, R.A., Moisan, L.J., Medina-Escobar, N., Neu, C., Cabral, A., and Parker, J.E. (2005). Arabidopsis SENESCENCE-ASSOCIATED GENE101 stabilizes and signals within an ENHANCED DISEASE SUSCEPTIBILITY1 complex in plant innate immunity. The Plant cell 17, 2601–2613.

Gantner, J., Ordon, J., Ilse, T., Kretschmer, C., Gruetzner, R., Lofke, C., Dagdas, Y., Burstenbinder, K., Marillonnet, S., and Stuttmann, J. (2018). Peripheral infrastructure vectors and an extended set of plant parts for the Modular Cloning system. PLoS ONE 13, e0197185.

Garcia, A.V., Blanvillain-Baufume, S., Huibers, R.P., Wiermer, M., Li, G., Gobbato, E., Rietz, S., and Parker, J.E. (2010). Balanced nuclear and cytoplasmic activities of EDS1 are required for a complete plant innate immune response. PLoS Pathog 6, e1000970.

Gietz, R.D., and Schiestl, R.H. (2007). Frozen competent yeast cells that can be transformed with high efficiency using the LiAc/SS carrier DNA/PEG method. Nat Protoc 2, 1–4.

Hecker, A., Wallmeroth, N., Peter, S., Blatt, M.R., Harter, K., and Grefen, C. (2015). Binary 2in1 Vectors Improve in Planta (Co)localization and Dynamic Protein Interaction Studies. Plant physiology 168, 776–787.

Heidrich, K., Wirthmueller, L., Tasset, C., Pouzet, C., Deslandes, L., and Parker, J.E. (2011). Arabidopsis EDS1 connects pathogen effector recognition to cell compartment-specific immune responses. Science (New York, N.Y 334, 1401–1404.

Hu, G., deHart, A.K., Li, Y., Ustach, C., Handley, V., Navarre, R., Hwang, C.F., Aegerter, B.J., Williamson, V.M., and Baker, B. (2005). EDS1 in tomato is required for resistance mediated by TIR-class R genes and the receptor-like R gene Ve. Plant J 42, 376–391.

Huh, S.U., Cevik, V., Ding, P., Duxbury, Z., Ma, Y., Tomlinson, L., Sarris, P.F., and Jones, J.D.G. (2017). Protein-protein interactions in the RPS4/RRS1 immune receptor complex. PLoS Pathog 13, e1006376.

Jacob, F., Vernaldi, S., and Maekawa, T. (2013). Evolution and Conservation of Plant NLR Functions. Front Immunol 4, 297.

Jones, J.D., and Dangl, J.L. (2006). The plant immune system. Nature 444, 323–329.

Jones, J.D., Vance, R.E., and Dangl, J.L. (2016). Intracellular innate immune surveillance devices in plants and animals. Science (New York, N.Y 354.

Kadota, Y., Shirasu, K., and Guerois, R. (2010). NLR sensors meet at the SGT1-HSP90 crossroad. Trends Biochem Sci 35, 199–207.

Khan, M., Subramaniam, R., and Desveaux, D. (2016). Of guards, decoys, baits and traps: pathogen perception in plants by type III effector sensors. Curr Opin Microbiol 29, 49–55.

Kim, S.H., Kwon, S.I., Saha, D., Anyanwu, N.C., and Gassmann, W. (2009). Resistance to the Pseudomonas syringae effector HopA1 is governed by the TIR-NBS-LRR protein RPS6 and is enhanced by mutations in SRFR1. Plant physiology 150, 1723–1732.

Kim, T.-H., Kunz, H.-H., Bhattacharjee, S., Hauser, F., Park, J., Engineer, C., Liu, A., Ha, T., Parker, J.E., Gassmann, W., and Schroeder, J.I. (2012). Natural Variation in Small Molecule-Induced TIR-NB-LRR Signaling Induces Root Growth Arrest via EDS1- and PAD4-Complexed R Protein VICTR in Arabidopsis. The Plant cell 24, 5177–5192.

Krasileva, K.V., Dahlbeck, D., and Staskawicz, B.J. (2010). Activation of an Arabidopsis resistance protein is specified by the in planta association of its leucine-rich repeat domain with the cognate oomycete effector. The Plant cell 22, 2444–2458.

Kushnirov, V.V. (2000). Rapid and reliable protein extraction from yeast. Yeast 16, 857–860.

Lipka, V., Dittgen, J., Bednarek, P., Bhat, R., Wiermer, M., Stein, M., Landtag, J., Brandt, W., Rosahl, S., Scheel, D., Llorente, F., Molina, A., Parker, J., Somerville, S., and Schulze-Lefert, P. (2005). Pre- and postinvasion defenses both contribute to nonhost resistance in Arabidopsis. Science (New York, N.Y 310, 1180–1183.

Liu, D., Shi, L., Han, C., Yu, J., Li, D., and Zhang, Y. (2012). Validation of reference genes for gene expression studies in virus-infected Nicotiana benthamiana using quantitative real-time PCR. PLoS ONE 7, e46451.

Logemann, E., Birkenbihl, R.P., Ulker, B., and Somssich, I.E. (2006). An improved method for preparing Agrobacterium cells that simplifies the Arabidopsis transformation protocol. Plant methods 2, 16.

Louis, J., Gobbato, E., Mondal, H.A., Feys, B.J., Parker, J.E., and Shah, J. (2012). Discrimination of Arabidopsis PAD4 activities in defense against green peach aphid and pathogens. Plant physiology 158, 1860–1872.

Macho, A.P., and Zipfel, C. (2014). Plant PRRs and the activation of innate immune signaling. Molecular cell 54, 263–272.

Macho, A.P., and Zipfel, C. (2015). Targeting of plant pattern recognition receptor-triggered immunity by bacterial type-III secretion system effectors. Curr Opin Microbiol 23, 14–22.

Maekawa, T., Kufer, T.A., and Schulze-Lefert, P. (2011a). NLR functions in plant and animal immune systems: so far and yet so close. Nat Immunol 12, 817–826.

Maekawa, T., Kracher, B., Vernaldi, S., Ver Loren van Themaat, E., and Schulze-Lefert, P. (2012). Conservation of NLR-triggered immunity across plant lineages. Proceedings of the National Academy of Sciences of the United States of America 109, 20119–20123.

Maekawa, T., Cheng, W., Spiridon, L.N., Toller, A., Lukasik, E., Saijo, Y., Liu, P., Shen, Q.H., Micluta, M.A., Somssich, I.E., Takken, F.L., Petrescu, A.J., Chai, J., and Schulze-Lefert, P. (2011b). Coiled-coil domain-dependent homodimerization of intracellular barley immune receptors defines a minimal functional module for triggering cell death. Cell Host Microbe 9, 187–199.

Mindrinos, M., Katagiri, F., Yu, G.-L., and Ausubel, F.M. (1994). The *A. thaliana* disease resistance gene *RPS2* encodes a protein containing a nucleotide-binding site and leucine-rich repeats. Cell 78, 1089–1099.

Mondragon-Palomino, M., Meyers, B.C., Michelmore, R.W., and Gaut, B.S. (2002). Patterns of positive selection in the complete NBS-LRR gene family of Arabidopsis thaliana. Genome research 12, 1305–1315.

Monteiro, F., and Nishimura, M.T. (2018). Structural, Functional, and Genomic Diversity of Plant NLR Proteins: An Evolved Resource for Rational Engineering of Plant Immunity. Annu Rev Phytopathol 56, 243–267.

Ordon, J., Gantner, J., Kemna, J., Schwalgun, L., Reschke, M., Streubel, J., Boch, J., and Stuttmann, J. (2017). Generation of chromosomal deletions in dicotyledonous plants employing a user-friendly genome editing toolkit. Plant J 89, 155–168.

Parker, J.E., Holub, E.B., Frost, L.N., Falk, A., Gunn, N.D., and Daniels, M.J. (1996). Characterization of eds1, a mutation in Arabidopsis suppressing resistance to Peronospora parasitica specified by several different RPP genes. The Plant cell 8, 2033–2046.

Peart, J.R., Cook, G., Feys, B.J., Parker, J.E., and Baulcombe, D.C. (2002). An *EDS1* orthologue is required for *N*-mediated resistance against tobacco mosaic virus. Plant J 29, 569–579.

Pegadaraju, V., Knepper, C., Reese, J., and Shah, J. (2005). Premature leaf senescence modulated by the Arabidopsis PHYTOALEXIN DEFICIENT4 gene is associated with defense against the phloem-feeding green peach aphid. Plant physiology 139, 1927–1934.

Pegadaraju, V., Louis, J., Singh, V., Reese, J.C., Bautor, J., Feys, B.J., Cook, G., Parker, J.E., and Shah, J. (2007). Phloem-based resistance to green peach aphid is controlled by Arabidopsis PHYTOALEXIN DEFICIENT4 without its signaling partner ENHANCED DISEASE SUSCEPTIBILITY1. Plant J 52, 332–341.

Pupko, T., Bell, R.E., Mayrose, I., Glaser, F., and Ben-Tal, N. (2002). Rate4Site: an algorithmic tool for the identification of functional regions in proteins by surface mapping of evolutionary determinants within their homologues. Bioinformatics 18 Suppl 1, S71–77.

Qi, T., Seong, K., Thomazella, D.P.T., Kim, J.R., Pham, J., Seo, E., Cho, M.J., Schultink, A., and Staskawicz, B.J. (2018). NRG1 functions downstream of EDS1 to regulate TIR-NLR-mediated plant immunity in Nicotiana benthamiana. Proceedings of the National Academy of Sciences of the United States of America.

Rietz, S., Stamm, A., Malonek, S., Wagner, S., Becker, D., Medina-Escobar, N., Vlot, A.C., Feys, B.J., Niefind, K., and Parker, J.E. (2011). Different roles of Enhanced Disease Susceptibility1 (EDS1) bound to and dissociated from Phytoalexin Deficient4 (PAD4) in Arabidopsis immunity. New Phytol 191, 107–119.

Rosli, H.G., Zheng, Y., Pombo, M.A., Zhong, S., Bombarely, A., Fei, Z., Collmer, A., and Martin, G.B. (2013). Transcriptomics-based screen for genes induced by flagellin and repressed by pathogen effectors identifies a cell wall-associated kinase involved in plant immunity. Genome biology 14, R139.

Sarris, P.F., Cevik, V., Dagdas, G., Jones, J.D., and Krasileva, K.V. (2016). Comparative analysis of plant immune receptor architectures uncovers host proteins likely targeted by pathogens. BMC Biol 14, 8.

Schon, M., Toller, A., Diezel, C., Roth, C., Westphal, L., Wiermer, M., and Somssich, I.E. (2013). Analyses of wrky18 wrky40 Plants Reveal Critical Roles of SA/EDS1 Signaling and Indole-Glucosinolate Biosynthesis for Golovinomyces orontii Resistance and a Loss-of Resistance Towards Pseudomonas syringae pv. tomato AvrRPS4. Molecular plant-microbe interactions : MPMI 26, 758–767.

Schultink, A., Qi, T., Lee, A., Steinbrenner, A.D., and Staskawicz, B. (2017). Roq1 mediates recognition of the Xanthomonas and Pseudomonas effector proteins XopQ and HopQ1. Plant J.

Shimada, T.L., Shimada, T., and Hara-Nishimura, I. (2010). A rapid and non-destructive screenable marker, FAST, for identifying transformed seeds of Arabidopsis thaliana. Plant J 61, 519–528.

Sinapidou, E., Williams, K., Nott, L., Bahkt, S., Tor, M., Crute, I., Bittner-Eddy, P., and Beynon, J. (2004). Two TIR:NB:LRR genes are required to specify resistance to Peronospora parasitica isolate Cala2 in Arabidopsis. Plant J 38, 898–909.

Sohn, K.H., Hughes, R.K., Piquerez, S.J., Jones, J.D.G., and Banfield, M.J. (2012). Distinct regions of the Pseudomonas syringae coiled-coil effector AvrRps4 are required for activation of immunity. Proceedings of the National Academy of Sciences of the United States of America 109, 16371–16376.

Solovyev, V.V. (2007). Statistical approaches in Eukaryotic gene prediction. In Handbook of Statistical Genetics.

Song, Y., DiMaio, F., Wang, R.Y., Kim, D., Miles, C., Brunette, T., Thompson, J., and Baker, D. (2013). High-resolution comparative modeling with RosettaCM. Structure 21, 1735–1742.

Spoel, S.H., and Dong, X. (2012). How do plants achieve immunity? Defence without specialized immune cells. Nat Rev Immunol 12, 89–100.

Steinbrenner, A.D., Goritschnig, S., and Staskawicz, B.J. (2015). Recognition and activation domains contribute to allele-specific responses of an Arabidopsis NLR receptor to an oomycete effector protein. PLoS Pathog 11, e1004665.

Stuttmann, J., Peine, N., Garcia, A.V., Wagner, C., Choudhury, S.R., Wang, Y., James, G.V., Griebel, T., Alcazar, R., Tsuda, K., Schneeberger, K., and Parker, J.E. (2016). Arabidopsis thaliana DM2h (R8) within the Landsberg RPP1-like Resistance Locus Underlies Three Different Cases of EDS1-Conditioned Autoimmunity. PLoS Genet 12, e1005990.

Swiderski, M.R., Birker, D., and Jones, J.D. (2009). The TIR domain of TIR-NB-LRR resistance proteins is a signaling domain involved in cell death induction. Mol Plant Microbe Interact 22, 157–165.

Takken, F.L.W., and Goverse, A. (2012). How to build a pathogen detector: structural basis of NB-LRR function. Current opinion in plant biology 15, 375–384.

Thieme, F., Koebnik, R., Bekel, T., Berger, C., Boch, J., Buttner, D., Caldana, C., Gaigalat, L., Goesmann, A., Kay, S., Kirchner, O., Lanz, C., Linke, B., McHardy, A.C., Meyer, F., Mittenhuber, G., Nies, D.H., Niesbach-Klosgen, U., Patschkowski, T., Ruckert, C., Rupp, O., Schneiker, S., Schuster, S.C., Vorholter, F.J., Weber, E., Puhler, A., Bonas, U., Bartels, D., and Kaiser, O. (2005). Insights into genome plasticity and pathogenicity of the plant pathogenic bacterium Xanthomonas campestris pv. vesicatoria revealed by the complete genome sequence. J Bacteriol 187, 7254–7266.

Thomas, W.J., Thireault, C.A., Kimbrel, J.A., and Chang, J.H. (2009). Recombineering and stable integration of the Pseudomonas syringae pv. syringae 61 hrp/hrc cluster into the genome of the soil bacterium Pseudomonas fluorescens Pf0-1. Plant Journal 60, 919–928.

Venugopal, S.C., Jeong, R.-D., Mandal, M.K., Zhu, S., Chandra-Shekara, A.C., Xia, Y., Hersh, M., Stromberg, A.J., Navarre, D., Kachroo, A., and Kachroo, P. (2009). Enhanced Disease Susceptibility 1 and Salicylic Acid Act Redundantly to Regulate Resistance Gene-Mediated Signaling. Plos Genetics 5.

Vlot, A.C., Dempsey, D.M.A., and Klessig, D.F. (2009). Salicylic Acid, a multifaceted hormone to combat disease. Annu Rev Phytopathol 47, 177–206.

Wagner, S., Stuttmann, J., Rietz, S., Guerois, R., Brunstein, E., Bautor, J., Niefind, K., and Parker, J.E. (2013). Structural basis for signaling by exclusive EDS1 heteromeric complexes with SAG101 or PAD4 in plant innate immunity. Cell Host Microbe 14, 619–630.

Weber, E., Engler, C., Gruetzner, R., Werner, S., and Marillonnet, S. (2011). A modular cloning system for standardized assembly of multigene constructs. PLoS ONE 6, e16765.

Wei, H.L., Zhang, W., and Collmer, A. (2018). Modular Study of the Type III Effector Repertoire in Pseudomonas syringae pv. tomato DC3000 Reveals a Matrix of Effector Interplay in Pathogenesis. Cell Rep 23, 1630–1638.

Wei, H.L., Chakravarthy, S., Mathieu, J., Helmann, T.C., Stodghill, P., Swingle, B., Martin, G.B., and Collmer, A. (2015). Pseudomonas syringae pv. tomato DC3000 Type III Secretion Effector Polymutants Reveal an Interplay between HopAD1 and AvrPtoB. Cell Host Microbe 17, 752–762.

Wiermer, M., Feys, B.J., and Parker, J.E. (2005). Plant immunity: the EDS1 regulatory node. Current opinion in plant biology 8, 383–389.

Williams, S.J., Sohn, K.H., Wan, L., Bernoux, M., Sarris, P.F., Segonzac, C., Ve, T., Ma, Y., Saucet, S.B., Ericsson, D.J., Casey, L.W., Lonhienne, T., Winzor, D.J., Zhang, X., Coerdt, A., Parker, J.E., Dodds, P.N., Kobe, B., and Jones, J.D. (2014). Structural basis for assembly and function of a heterodimeric plant immune receptor. Science (New York, N.Y 344, 299–303.

Wirthmueller, L., Zhang, Y., Jones, J.D., and Parker, J.E. (2007). Nuclear accumulation of the Arabidopsis immune receptor RPS4 is necessary for triggering EDS1-dependent defense. Curr Biol 17, 2023–2029.

Wu, Z., Li, M., Dong, O.X., Xia, S., Liang, W., Bao, Y., Wasteneys, G., and Li, X. (2018). Differential regulation of TNL-mediated immune signaling by redundant helper CNLs. New Phytol.

Yu, X., Feng, B., He, P., and Shan, L. (2017). From Chaos to Harmony: Responses and Signaling upon Microbial Pattern Recognition. Annu Rev Phytopathol 55, 109–137.

Zembek, P., Danilecka, A., Hoser, R., Eschen-Lippold, L., Benicka, M., Grech-Baran, M., Rymaszewski, W., Barymow-Filoniuk, I., Morgiewicz, K., Kwiatkowski, J., Piechocki, M., Poznanski, J., Lee, J., Hennig, J., and Krzymowska, M. (2018). Two Strategies of Pseudomonas syringae to Avoid Recognition of the HopQ1 Effector in Nicotiana Species. Front Plant Sci 9, 978.

Zhang, M., Kadota, Y., Prodromou, C., Shirasu, K., and Pearl, L.H. (2010). Structural basis for assembly of Hsp90-Sgt1-CHORD protein complexes: implications for chaperoning of NLR innate immunity receptors. Molecular cell 39, 269–281.

Zimmermann, L., Stephens, A., Nam, S.Z., Rau, D., Kubler, J., Lozajic, M., Gabler, F., Soding, J., Lupas, A.N., and Alva, V. (2018). A Completely Reimplemented MPI Bioinformatics Toolkit with a New HHpred Server at its Core. Journal of molecular biology 430, 2237–2243.

Zouine, M., Maza, E., Djari, A., Lauvernier, M., Frasse, P., Smouni, A., Pirrello, J., and Bouzayen, M. (2017). TomExpress, a unified tomato RNA-Seq platform for visualization of expression data, clustering and correlation networks. Plant J 92, 727–735.

